# Pentachloropseudilin disrupts multiple steps of the *Toxoplasma gondii* lytic cycle, consistent with predicted multi-myosin engagement

**DOI:** 10.64898/2026.09.30.755571

**Authors:** Romane Agresti, Faraj Gtat, Guillaume Drumont, Bastien Touquet, Christopher Swale, Isabelle Tardieux

## Abstract

The pharmacological targeting of apicomplexan myosins has emerged as a validated antiparasitic strategy, with selective inhibitors of MyoA demonstrating activity against parasite motility and invasion. However, apicomplexan parasites express multiple unconventional myosins with essential functions throughout their life cycles that remain unexplored pharmacologically. Here, we investigated pentachloropseudilin (PCIP), a reversible allosteric inhibitor characterized against class-I myosins, although canonical class-I myosins are absent from Apicomplexa. Structural modeling predicted PCIP-potentially compatible allosteric pockets in several parasite myosins, including MyoA, MyoF and MyoJ, suggesting interference with multiple myosin-dependent processes. To specifically resolve PCIP-induced phenotypes across the tachyzoite lytic cycle, we combined complementary high-content and high-resolution approaches, including real-time live-cell imaging, quantitative assays and ultrastructure expansion microscopy. This identified distinct steps at which parasite progression was impaired. PCIP rapidly and reversibly inhibited gliding motility and host-cell invasion in a dose-dependent manner within the submicromolar range. A pronounced effect was observed during intracellular development, where PCIP inhibited proliferation and arrested parasites near the onset of daughter-cell budding. Live imaging and expansion microscopy revealed failure to assemble daughter inner membrane complex scaffolds and subpellicular microtubule arrays, and to replicate apicoplast, consistent with disruption of MyoF-dependent processes. This arrest was not simply cytostatic, as even short PCIP exposure prevented a substantial proportion of intracellular tachyzoites from resuming replication following drug removal. At lower concentrations permitting daughter-cell formation, progeny displayed defective final maturation and frequently lost plasma-membrane integrity within an intact parasitophorous vacuole, preventing egress consistent with interference with MyoJ-dependent processes. Together, these findings reveal multiple points of PCIP-induced failure throughout the *T. gondii* lytic cycle and associate distinct phenotypes with MyoA-, MyoF- and MyoJ-dependent functions. They establish PCIP as a chemical probe for dissecting apicomplexan myosin-dependent processes and support multi-myosin interference as a promising antiparasitic strategy.

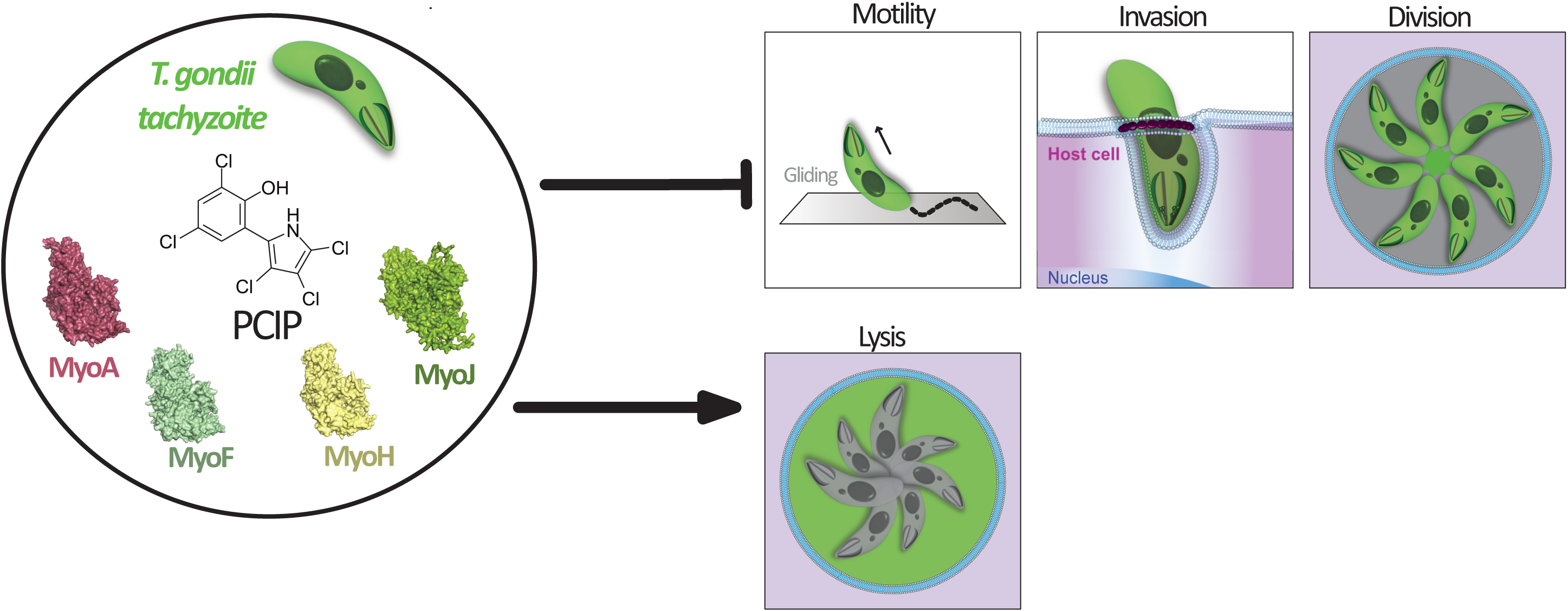

## Introduction

The Apicomplexa phylum comprises thousands of obligate intracellular protozoan parasites infecting virtually all animal phyla. Descended from free-living, photosynthetic ancestors, apicomplexans have diversified into one of the most widespread groups of animal parasites (<u>Janouškovec et al., 2015</u>). They are responsible for major diseases in humans and animals, including malaria due to *Plasmodium spp*. (<u>WHO, 2025</u>), cryptosporidiosis associated with *Cryptosporidium spp*. (<u>Sow et al., 2018</u>), and economically important infections by *Eimeria* and *Babesia spp*. (<u>Fayer, 1980</u>; <u>Blake et al., 2020</u>; <u>Kumar et al., 2021</u>). Among apicomplexans, *Toxoplasma gondii* has become one of the most extensively studied species because of its medical and veterinary importance and experimental tractability.

*T. gondii*, the sole recognized species of the genus *Toxoplasma*, is one of the world’s most prevalent protozoan parasites, owing to its broad host range, cosmopolitan distribution, and capacity to establish lifelong infections (<u>Hill & Dubey, 2002</u>). Approximately one-third of the global human population has been exposed to the parasite (<u>Montoya & Liesenfeld, 2004</u>; <u>Robert-Gangneux & Dardé, 2012</u>). Although infection is usually asymptomatic in immunocompetent individuals, persistent tissue cysts can reactivate following loss of immune control and cause life-threatening disease. Primary infection during pregnancy may also result in severe congenital toxoplasmosis, while ocular toxoplasmosis is a leading cause of infectious posterior uveitis worldwide (<u>Kalogeropoulos et al., 2022</u>).

Despite their diverse host ranges and life cycles, apicomplexans share the ability to actively invade host cells. Specialized motile stages, collectively termed “zoites”, use their pronounced apico-basal polarity and actomyosin-based motor machineries, known as glideosomes, to move and invade cells. Because invasion is essential for parasite propagation, the myosin system represents an attractive therapeutic target. This concept has gained pharmacological support through parasite-selective inhibitors of myosin A (MyoA) (<u>Kelsen et al., 2023</u>; <u>Du Toit, 2023</u>). KNX-002 inhibits *T. gondii* MyoA and tachyzoite motility, while KNX-115 targets *Plasmodium falciparum* MyoA with nanomolar potency (<u>Moussaoui et al., 2023</u>). More recently, <u>Trivedi et al., (2026)</u> demonstrated KNX-115 activity against MyoA across *Plasmodium*, *Cryptosporidium*, *Toxoplasma*, and *Eimeria* and resolved the PfMyoA–KNX-115 structure. Independently, <u>Snyder et al. (2026)</u> identified UCB-9721 as a potent TgMyoA inhibitor with activity against *Plasmodium*, *Cryptosporidium*, and *Babesia* and characterized its binding pocket. Together, these studies establish MyoA as a conserved and pharmacologically tractable target across Apicomplexa.

Beyond MyoA, apicomplexans encode a diverse repertoire of unconventional myosins assigned to several lineage-enriched classes, although classifications vary between phylogenetic frameworks (<u>Foth et al., 2006</u>; <u>Mueller et al., 2017</u>). These motors retain the conserved ATPase domain and support processes extending beyond gliding motility, including replication, cytokinesis, organelle inheritance, vesicular trafficking, and egress. MyoF, for example, is conserved across Apicomplexa and regulates actin organization, organelle positioning, apicoplast inheritance, and vesicular trafficking in *T. gondii* (<u>Jacot et al., 2013</u>; <u>Heaslip et al., 2016</u>; <u>Carmeille et al., 2021</u>; <u>Kellermeier and Heaslip, 2024</u>; <u>Sladewski and Heaslip, 2026</u>), while PfMyoF is associated with blood-stage growth (<u>Holmes et al., 2026</u>). TgMyoJ contributes to tachyzoite maturation through actin-dependent basal-complex constriction (<u>Frénal et al., 2017</u>). These motors therefore represent additional potential pharmacological targets.

More broadly, conservation of the ATPase motor domain raises the possibility that a single compound could target multiple parasite myosins and disrupt several essential processes simultaneously. Although apicomplexans lack canonical class-I myosins, pentachloropseudilin (PCIP) is a hydrophobic, reversible class-I myosin inhibitor that binds an allosteric pocket within the motor domain containing both conserved and isoform-specific residues (<u>Chinthalapudi et al., 2011</u>). We therefore asked whether structurally compatible allosteric pockets are retained across divergent apicomplexan myosins, potentially allowing PCIP to engage multiple parasite motors despite the absence of canonical class-I myosins.

To address this hypothesis, we used *T. gondii* as a model and compared AlphaFold 3 predictions of myosin/PCIP/ADP complexes for four myosins expressed in motile, rapidly replicating tachyzoites (MyoH, MyoA, MyoF, and MyoJ) with evolutionarily distant class-I myosins. Guided by these predictions, we combined live-cell imaging, quantitative image analysis, high-content phenotypic profiling, and expansion microscopy to assess PCIP effects on gliding motility, invasion, intracellular replication, and egress. Our findings show that PCIP broadly disrupts the *T. gondii* tachyzoite lytic cycle, supporting its use as a chemical probe of myosin-dependent processes and as a starting point for anti-*Toxoplasma* strategies targeting multiple motors.

## Methods

### Buffer, media and pharmacological inhibitors

– Complete medium: Dulbecco’s Modified Eagle Medium (DMEM; Gibco-Life Technologies) supplemented with 10 mM HEPES, 2 mM GlutaMAX, 10% heat-inactivated fetal calf serum (FCS), penicillin (100 U/mL), and streptomycin (100 µg/mL)
– MB: Motility Buffer (1× HBSS, 1% Foetal calf serum (FCS), and 1.6 mM CaCl₂)
– WB: Washing Buffer (1× HBSS, 0.2% FCS)
– LLIM: Long Live Imaging Medium: Fluorobright supplemented with 10 mM HEPES, 2mM GlutaMAX, 10% heat-inactivated fetal bovine serum (FBS), penicillin (100 U/mL), and streptomycin (100 µg/mL) at 37°C and 5% CO₂.
– PHEM 1X: 60 mM Piperazine-N,N′-bis(2-ethanesulfonic acid), 25 mM 4-(2-Hydroxyethyl)-1-piperazineethanesulfonic acid, 10 mM Ethylene glycol-bis(2-aminoethylether)-N,N,N′,N′-tetraacetic acid, 2 mM Magnesium chloride.
– PCIP powder was prepared as 10 mM stock solution in DMSO, aliquoted in 20 μL and kept at –80°C. Aliquots were discarded after use.
– A23187 powder was prepared as 2 mM stock solution in DMSO and aliquoted in 50 μL and kept at –20°C. It was reused for several assays with similar efficacy
– U-ExM monomer solution: 10% acrylamide, 19% sodium acrylate, 0.1% N,N′-(1,2-dihydroxyethylene)bisacrylamid (DHEBA) in PBS, stored as 100 μL aliquots at -20°C
– U-ExM denaturation buffer: 200 mM sodium dodecyl sulphate (SDS), 200 mM NaCl, 50 mM Tris base, adjusted to pH 6.8 in ddH_2_O stored as aliquots at –20°C.

### Structural modeling

Full-length myosin proteins were modelled using the AlphaFold Server (https://alphafoldserver.com/), using default parameters. Amino acid sequences of the proteins were taken from UniProt (O00159 · MYO1C_HUMAN; O00934 · MYOA_TOXGO; Q286V9 · MYOF_TOXGO; A0A7J6K9H8 · MYOH_TOXGO; A0A7J6K1W9 · MYOJ_TOXGO).

AlphaFold3 (AF3; Abramson et al., Nature, 2024) Myo/PCIP/ADP co-complex modelling was performed using standard parameters on a workstation, using AF3 model access provided to CS by Google DeepMind. Calculations were run iteratively on the following GPUs: an A5000, an RTX 4090, and an RTX 4080. The PCIP SMILES string (OC1=C(C2=C(Cl)C(Cl)=C(Cl)N2)C=C(Cl)C=C1Cl) was used to define the ligand for co-folding. Modelling was performed with and without ADP.The crystal structure of KNX-002-bound P. falciparum MyoA was also used as an experimental reference (PDB: 8CDQ; <u>Moussaoui et al., 2023</u>). All visual representations were generated using the PyMOL Molecular Graphics System, Version 3.0, Schrödinger, LLC.

### Human and Parasite cell Production

Human foreskin fibroblasts (HFFs), A549 human lung adenocarcinoma epithelial cells, and U2OS human osteosarcoma cells were maintained in complete DMEM, at 37 °C in a humidified atmosphere containing 5% CO₂. Cells were cultured in T25 or T75 tissue-culture flasks and routinely passaged before reaching full confluence. For maintenance, cells were detached using trypsin-EDTA and serially diluted into fresh complete DMEM before being transferred to new culture flasks.

All *Toxoplasma gondii* strains used in this study were genetically engineered derivatives of the type I RHΔku80 background, which facilitates targeted homologous recombination through disruption of the non-homologous end-joining pathway (<u>Huynh and Carruthers, 2009</u>). Where applicable, endogenous tagging was performed using ligation-independent cloning (LIC)-based approaches. The strains used in this study were RHΔku80 UPRT::NLuc-P2A-EmGFP (<u>Bellini et al., 2020</u>) RHΔku80 RON2-3HA DHFR, RHΔku80 RON2mC-HX (<u>Bichet et al., 2014</u>, and RHΔku80 IMC12mC-BCC7Em, RHΔku80 P12mC-BCC7Em; (<u>Vigetti et al., 2022</u>). *T. gondii* tachyzoites were maintained by serial passage in confluent monolayers of HFFs under standard culture conditions (37°C, 5% CO₂).

### Real-time live-cell imaging assays

All assays were performed using the confocal inverted microscope EclipseTi (Nikon France Instruments, Champigny sur Marne France) equipped with the thermo and gaz controllers to achieve 37°C and 5% CO_2_ atmosphere (Tokai Hit, Japan). The microscope was equipped with a piezo element and PrismBSI express camera (Photometrics), a live SR module of super resolution (Gataca), and a CSU X1 spinning disk (Yokogawa). Image capure was performed using the Metamorph software (Universal Imaging Corporation, Roper Scientific, Lisses, France).

#### Tachyzoite motility assays

For tachyzoite 2D motility, live-cell imaging was performed with parasites settled on cleaned 18-mm glass coverslips previously equilibrated in MB. Tachyzoites were collected following spontaneous egress from HFF monolayers. The suspension was first centrifuged at 200 *g* for 2 min and filtered through a 5-µm PluriStrainer to remove cellular debris, while the free tachyzoites containing were collected by centrifugation at 800 *g* for 5 min in WB, using a volume corresponding to five times the initial suspension volume. The tachyzoite-enriched pellet was resuspended in MB supplemented with either 0.01% DMSO (vehicle, control) or PCIP pharmacological inhibit at final concentrations of 0.1, 0.5, or 1 µM. About 4-5 10⁵ tachyzoites were loaded into a final volume of 200 µL and deposited onto the 18-mm coverslip in a magnetic microscope chamber. The chamber was incubated for 2 min at 37°C and subsequently centrifuged in a swinging-bucket rotor at 200 *g* for 2 min to facilitate parasite deposition onto the coverslip. The chamber was then transferred to the microscope. After 1-2 min to stabilize the setting, tachyzoite behavior was recorded for 5.5 min at a rate of two frames per second (2 fps) using a 40x Pan Fluor (1.30 NA, Nikon) oil immersion objective.

To assess the reversibility of the effects of PCIP on tachyzoite motility, the parasites were exposed to PCIP or the DMSO vehicle control for 5 min and subsequently washed with WB, using a volume corresponding to at least 70 times the sample volume to remove the compound. Following centrifugation at 800 *g* for 5 min, the parasite containing pellet was resuspended in a 200 µL of MB and incubated for 30 min at 37°C. Parasites were then loaded into the magnetic microscope chamber, and CaCl₂ was added to a final concentration of 1.6 mM. Samples were subsequently handled as described above and tachyzoite behavior recorded for 5.5 min at a rate of 2 frames per second (fps) using 40x Pan Fluor (1.30 NA, Nikon) oil immersion objective.

To assess the effect of calcium ionophore treatment on PCIP-treated parasites, tachyzoites were exposed to PCIP or the DMSO vehicle control as described above and subsequently loaded into the magnetic microscope chamber. Tachyzoite behavior was recorded for 6 min at a rate of 2 fps. After the acquisition of the first 100 frames, calcium ionophore A23187 was added to a final concentration of 1 µM, and imaging was continued immediately thereafter. For assessment of the ionophore response, parasite behavior was subsequently analyzed during the period of calcium ionophore treatment.

#### Host cell invasion assays

To characterize tachyzoite behavior during host cell invasion, parasites expressing RON2-mC were deposited onto subconfluent U2OS cells stably expressing the GFP-GPI chimeric protein as a plasma membrane reporter (<u>Bichet et al., 2014</u>). Parasites were deposited onto the cells and let to settle for a couple of minutes, in the presence of vehicle or PCIP at concentrations of 0.1, 0.5, or 1 μM. Tachyzoite behavior was recorded for 17 min using three channels (λex 488 nm, λex 561 nm, and transmitted light) at a frame rate of 1.5 frames per second (fps), using a 60× Plan Fluor oil-immersion objective (NA 1.30, Nikon). This setup enabled monitoring of RON complex release at the target cell cortex and its organization into a torus, concomitant with the formation of a nascent PV.

#### Intracellular tachyzoite growth assays

To monitor the effect of PCIP on intracellular tachyzoite growth, HFFs were seeded on glass coverslips in 24-well plates (∼3.5 10^4^ cells/well) and cultured for 48 h to confluence before the infection with fluorescent tachyzoites (expressing either nLuc-GFP, IMC12-mC-BCC7Em, or P12mC-BCC7Em)(<u>Bichet et al., 2014</u>; <u>Pavlou et al., 2018</u>;<u>Vigetti et al., 2022</u>). Parasite invasion was monitored at 15-min intervals to ensure an appropriate level of infection. Next, extracellular tachyzoites were removed by washing with HBSS, and infected cells were maintained in live-cell imaging medium (LLIM) at 37°C and 5% CO₂. Between 1 and 2 h post-invasion, the medium was replaced with 5 mL of pre-warmed LLIM containing the appropriate concentration of PCIP or vehicle control. The MatTek dish was then mounted in the microscope chamber and maintained under controlled temperature and CO₂ conditions throughout image acquisition. Imaging parameters, including laser power and exposure time, were optimized to minimize phototoxicity during prolonged acquisition. Bright-field and fluorescence images corresponding to the expressed reporters were acquired every 4 min for up to 30 h, allowing continuous monitoring of intracellular parasite development and replication.

### Quantitative analysis of *T. gondii* tachyzoite cell invasiveness

HFFs were seeded on glass coverslips in 24-well plates (∼3.5 10^4^ cells/well) and cultured for 48 h to confluence. GFP-expressing tachyzoites, prepared as for motility assays, were pre-incubated for 5 min at 37°C and 5% CO₂ with vehicle or PCIP (0.05–10 µM) in 1 mL complete medium, then added to HFF monolayers at an MOI of ∼3:1. Invasion proceeded for up to 2 h and was monitored using an inverted microscope (Axio Vert.A1 FL, Zeiss), after which extracellular parasites were removed by PBS washing.

Samples were fixed with freshly prepared 3.2% PFA in PHEM buffer for 25 min at room temperature (RT) and blocked with 2% (w/v) BSA in PBS for 20 min. To distinguish extracellular from intracellular parasites, non-permeabilized samples were immunostained with anti-SAG1 antibody for 25 min, followed by goat anti-mouse Alexa Fluor 594-conjugated secondary antibody for 25 min, at RT. Nuclei were counterstained with Hoechst (1 µg/mL in PBS) for 10 min, and coverslips were washed with PBS and mounted in Mowiol.

For each of three independent experiments, 50 images per condition were acquired using an ApoTome AxioImager M2 fluorescence microscope (Zeiss) with a 40× oil-immersion objective. Invasiveness was quantified in Fiji using a custom macro developed with AI-assisted coding. The macro segments host-cell nuclei (Hoechst 33342) and GFP-positive parasites, determines parasite colocalization with the red surface-antigen marker, and can process up to 1,400 fields in <12 h. Detection thresholds were fixed across all images and conditions. Counting accuracy was validated against manual counts on randomly selected images using a quality-control macro (Fig. Sup. 3a). Complete workflow, including both macros, representative raw and processed images, and detailed invasion and quality-control results, is available at <u>the Tardieux-lab PCIP invasion macros repository</u>.

An additional invasion assay, performed in triplicate or quadruplicate, used HFFs seeded at 2 10^4^ cells/well in clear-bottom 96-well plates 48 h before infection and an MOI of 10:1. Samples were processed using the same immunolabeling protocol and imaged with an Olympus ScanR high-content screening microscope using a 20× objective. Thirty non-overlapping fields per well were acquired in three fluorescence channels (λex 405, 520, and 610 nm), as described by <u>Touquet et al. (2018)</u>.

For IC₅₀ determination, invasion values from each experiment were normalized to the vehicle control (100%) to account for differences in baseline invasion. Normalized responses were plotted against log₁₀[PCIP] (µM) and fitted in GraphPad Prism using a four-parameter log(inhibitor) *vs.* response model with variable slope. The top plateau was constrained to 100%, whereas the bottom was unconstrained; vehicle controls defined the top plateau but were excluded from the regression. IC₅₀ values were derived from fitted curves and are reported in µM as mean ± SD.

### Quantitative analysis of *T. gondii* tachyzoite growth and reversibility assays

HFF monolayers were prepared as described above for the invasion assays and infection was conducted using GFP-expressing tachyzoites handled as described for the host cell invasion assay. Infection levels were monitored under the inverted microscope (Axio Vert.A1 FL, Zeiss) to minimize the occurrence of host cells containing multiple independently established parasitophorous vacuoles (PVs). Following invasion, extracellular parasites were removed by washing with PBS. Infected cells were then incubated in complete medium for 15 min in the culture incubator, after which the medium was replaced with fresh complete medium containing either vehicle or PCIP at 0.1 μM, 0.5 μM and 1 μM in a final volume of 1 mL per well.

For the growth assay, cell samples were fixed at 22 h post-invasion with PFA 3.2% in PBS for 25 min, and stained with Hoechst (1 mg/mL in PBS) for 10 min at room temperature followed by PBS washes and mounting in Mowiol. For the growth reversibility assay, two treatment duration were tested. Infected cells were treated with 0.5 μM PCIP for either 8 or 20 h, after which the compound was removed by extensive washing with complete medium. Samples were subsequently incubated for an additional 24 h at 37°C under 5% CO₂ to allow parasite growth following PCIP removal. An additional control was included in which HFFs were infected at the time of PCIP removal to assess parasite growth over a 24-h period under the same experimental conditions.

For all assays, images were acquired using an ApoTome AxioImager M2 upright fluorescence microscope (Zeiss) equipped with a 40× oil-immersion objective (Zeiss) and subsequently processed using Fiji. Parasite growth was quantified by determining the number of tachyzoite progeny per PV. Quantification was performed across three independent experiments, each performed in duplicates with a minimum of 250 PVs analyzed per condition and assay. Recovery of parasite growth following PCIP removal was quantified by determining the proportion of PVs containing 1, 2, or more than 4 parasites per PVs. A minimum of 1,000 PVs was analyzed across three independent experiments.

### Quantitative analysis of the late-stage effect of PCIP on growing parasitophorous vacuoles

HFF monolayers were prepared as described above for the invasion and intracellular growth assays, and GFP-expressing tachyzoites were prepared as described for the motility assays. Infection levels were monitored to minimize the occurrence of host cells containing multiple independently established parasitophorous vacuoles (PVs). Following invasion, extracellular parasites were removed by washing with PBS. Infected cells were then incubated in complete medium for 15 min before treatment with medium containing either vehicle (DMSO), 0.1 μM PCIP, or 2.5 μM PCIP. Cells were subsequently cultured for 30–36 h at 37°C in a humidified atmosphere containing 5% CO₂. Cells were fixed with 3.2% PFA as described above, and nuclei were counterstained with Hoechst (1 mg/mL in PBS) for10 min. Coverslips were then washed and mounted in Mowiol. Images were acquired using a Nikon Eclipse Ti microscope with the Live-SR module and using the 60× Plan-neofluor oil immersion objective (1.46 NA, Zeiss). Image data were subsequently processed and analyzed using Fiji. Quantitative analysis of tachyzoite fate (growth or lysis) was performed by analyzing PVs from 10 images randomly taken per condition.

### Cell Cytotoxicity SYTOX-base assay

To assess the cytotoxicity of PCIP, HFF and A549 cells were seeded at a density of approximately 3.5 10⁴ cells per well and cultured for 48 h to obtain confluent monolayers. PCIP was diluted in complete medium to obtain increasing concentrations ranging from 0.05 to 80 μM. Untreated cells containing the same final concentration of vehicle were included as the negative viability control. Cells were cultured for 24 h. As a positive control for cell death, cells were incubated with 0.1% Triton X-100 for 10 min before staining. Cell death was quantified using SYTOX Green staining in combination with Hoechst 33342 nuclear staining. Cells were first incubated in complete medium containing 1 µg/mL Hoechst 33342 for 10 min at 37°C in a humidified atmosphere containing 5% CO₂. A 10× SYTOX Green solution was then added to achieve a final concentration of 60 nM, followed by an additional 5-min incubation. Excess Hoechst 33342 and SYTOX Green were carefully removed by gently replacing the culture medium to minimize cell detachment. Cells were subsequently fixed with 2% PFA for 30 min to preserve fluorescence signals and cellular morphology for automated image acquisition.

Images from 30 non-overlapping fields per well were automatically acquired using an Olympus ScanR High-Content Screening Station equipped with a 20× objective, using excitation wavelengths of 350 nm for Hoechst 33342 and 488 nm for SYTOX Green. Image processing and quantitative analysis were performed using Olympus ScanR Analysis software as described in <u>Touquet et al. (2018)</u>. Hoechst-positive nuclei were segmented to determine the total number of cells per well, whereas SYTOX Green-positive nuclei were independently segmented to determine the number of dead or membrane-compromised cells. The proportion of dead cells was normalized to the corresponding DMSO vehicle control. Quantitative analyses were performed using a minimum of two independent experiments both for HFF and A549 cells, and in duplicates for all PCIP concentrations.

### Ultrastructure expansion microscopy

Ultrastructure expansion microscopy (U-ExM) was performed using a modified protocol adapted from <u>Louvel et al. (2023)</u>, with N,N′-(1,2-dihydroxyethylene)bisacrylamide (DHEBA) replacing N,N′-methylenebisacrylamide as the hydrogel cross-linker. HFF monolayers grown on 12-mm coverslips were infected with *T. gondii* tachyzoites for 1 h, washed with PBS, and cultured for an additional 24 h at 37°C and 5% CO₂. Samples were anchored overnight at 37°C in 2% acrylamide/1.4% formaldehyde in PBS and placed in gelation chambers producing ∼170-µm-thick gels. Samples were covered with U-ExM monomer solution prepared in PBS, with APS and TEMED added immediately before gelation, incubated on ice for 15 min and polymerized at 37°C for 45 min. Gels were detached from coverslips in U-ExM denaturation buffer under gentle agitation and denatured in fresh buffer for 90 min at 85°C. Gels were then expanded in ddH₂O using at least three successive 20–30-min water exchanges at room temperature.

Expanded gels were cut to size, transferred to PBS to induce shrinkage, and incubated overnight at 4°C with mouse anti-IMC1 (1:400; gift from G. Ward, University of Vermont, USA) or anti-acetylated α-tubulin (1:500; Cell Signaling Technology), or rabbit anti-Cpn60 (1:600) in PBS/2% BSA. After extensive washing in PBS/0.1% Tween 20, gels were incubated for 2.5 h at 37°C with appropriated pre-absorbed goat anti-mouse or anti-rabbit IgG-ATTO 647N (1:600; Rockland), followed by ATTO 488 NHS ester (5 µg/mL; Sigma) for 1 h at room temperature. Gels were washed three times in PBS/0.1% Tween 20 and re-expanded through successive ddH₂O washes, with propidium iodide (1 µg/mL) included in the first water bath, until maximal expansion.

Fully expanded gels were mounted on poly-D-lysine-coated coverslips and imaged using the Nikon Eclipse Ti microscope described above, equipped with a Plan Fluor 40× oil-immersion objective (NA 1.3). Z-stacks were acquired at 0.2-µm intervals and processed in Fiji. The macroscopic linear expansion factor, calculated as the ratio of expanded to unexpanded gel dimensions, ranged from 4.5 to 5 across experiments.

### Image and Video analysis

Image and video analyses were performed using MetaMorph and Fiji software, unless otherwise specified. All images shown are representative of independent replicates.

For motility assays, individual tachyzoites were manually analyzed throughout the recordings and classified according to their observed motility phenotype, including active movement, specific movement patterns, or complete immobility. Time-lapse sequences were extracted from the full-length recordings of motility, invasion, and intracellular growth assays and annotated to highlight features or events of interest. For motility recordings, the Manual Tracking plugin in Fiji was used to track parasite displacement and reconstruct individual trajectories.

For static imaging of intracellular tachyzoite fate, including growth, growth arrest, and death, using epifluorescence, confocal, or expansion microscopy, maximum-intensity projections of fluorescence z-stacks are shown in the corresponding figure panels and annotated to highlight features of interest.

### Statistical analysis

Data were analyzed using GraphPad Prism (version 8.0.2). Statistical tests were selected according to the experimental design and type of data analyzed.

For motility assays, the behavior of each parasite was assessed throughout the recording sequence and classified according to the type of movement observed. The proportion of parasites exhibiting each type of movement was calculated for each experiment and condition and expressed as a percentage of motility. Three independent experiments were performed. A repeated-measures one-way ANOVA with Geisser–Greenhouse correction was performed across all experimental conditions. Data are presented as the mean ± SD of three independent experiments (*n* = 3). Statistical significance was assessed using a significance threshold of α = 0.05. Statistical significance is indicated as \*\*\*\**p* < 0.0001.

For invasion assays, the number of intracellular parasites per 100 host cells was determined for each experiment and condition. As each experimental value represented a mean derived from several thousand cells, a Gaussian approximation was considered appropriate. Conditions were treated as paired within each experiment, as measurements were obtained from the same parasite sample and drug aliquot and were subjected to the same fixation, washing, and staining procedures. A repeated-measures one-way ANOVA with Geisser–Greenhouse correction was performed on the six conditions represented in all three independent experiments. A test for linear trend was used to assess whether parasite invasion decreased with increasing PCIP concentration. The two additional concentrations (0.4 µM and 10 µM PCIP), for which data were available from only two of the three experiments, were analyzed using a mixed-effects model fitted by restricted maximum likelihood (REML), allowing the inclusion of incomplete datasets. Uncorrected Fisher’s LSD tests were used for pairwise comparisons between individual PCIP concentrations and the control. Data are presented as the mean of three independent experiments (*n* = 3), except for the 0.4 and 10 µM PCIP conditions (*n* = 2), with error bars as indicated in the figure legends. Statistical significance was assessed using a significance threshold of α = 0.05.

For the intracellular growth comparison, we compared first the distribution of parasites per PV across all conditions by two-way repeated-measures ANOVA, with the number of parasites per vacuole (1, 2, 4, 8, 16) and PCIP concentration as the two factors, matched by experiment on both of them; sphericity was not assumed (Geisser–Greenhouse correction). Because the treatment × category interaction was significant, we could perform pairwise comparisons as separate two-way repeated-measures ANOVAs, in which the interaction term tests whether the two distributions differ for a given pair for independent experiments (*n* = 3).

For late-stage maturation (*n* = 1) and delayed drug-release assays (*n* = 3), a PV was scored as imploded when the enclosed parasites had lysed and released GFP into an otherwise intact PV. Imploded and non-imploded vacuoles were treated as categorical outcomes, and their proportions were compared between conditions using Fisher’s exact test. A one-sided test was used based on the *a priori* hypothesis that PCIP treatment would increase the frequency of parasite lysis. Data are presented as percentages, with the corresponding absolute counts provided in the figure legends.

## Results

### Structural modelling of PCIP placement

We first generated AlphaFold3 (AF3) models of human Myo1c and the *T. gondii* myosins MyoA, MyoF, MyoH, and MyoJ to assess the structural conservation of their motor domains. We then compared these models with the crystal structure of KNX-bound *P. falciparum* MyoA (PfMyoA; PDB: 8CDQ), revealing broad conservation of the motor-domain architecture across human and parasite myosins. Given the high degree of MyoA conservation among apicomplexan parasites (with TM scores from 0.76 to 0.89), the PfMyoA crystal structure also provided a relevant structural reference for the binding site of the known MyoA inhibitor KNX (Fig. 1a, b), which isfound in the immediate vicinity of the ATP binding site and coodinates a water molecule, together with the γ-thiophosphate group of ATPγS.

**Figure 1.**
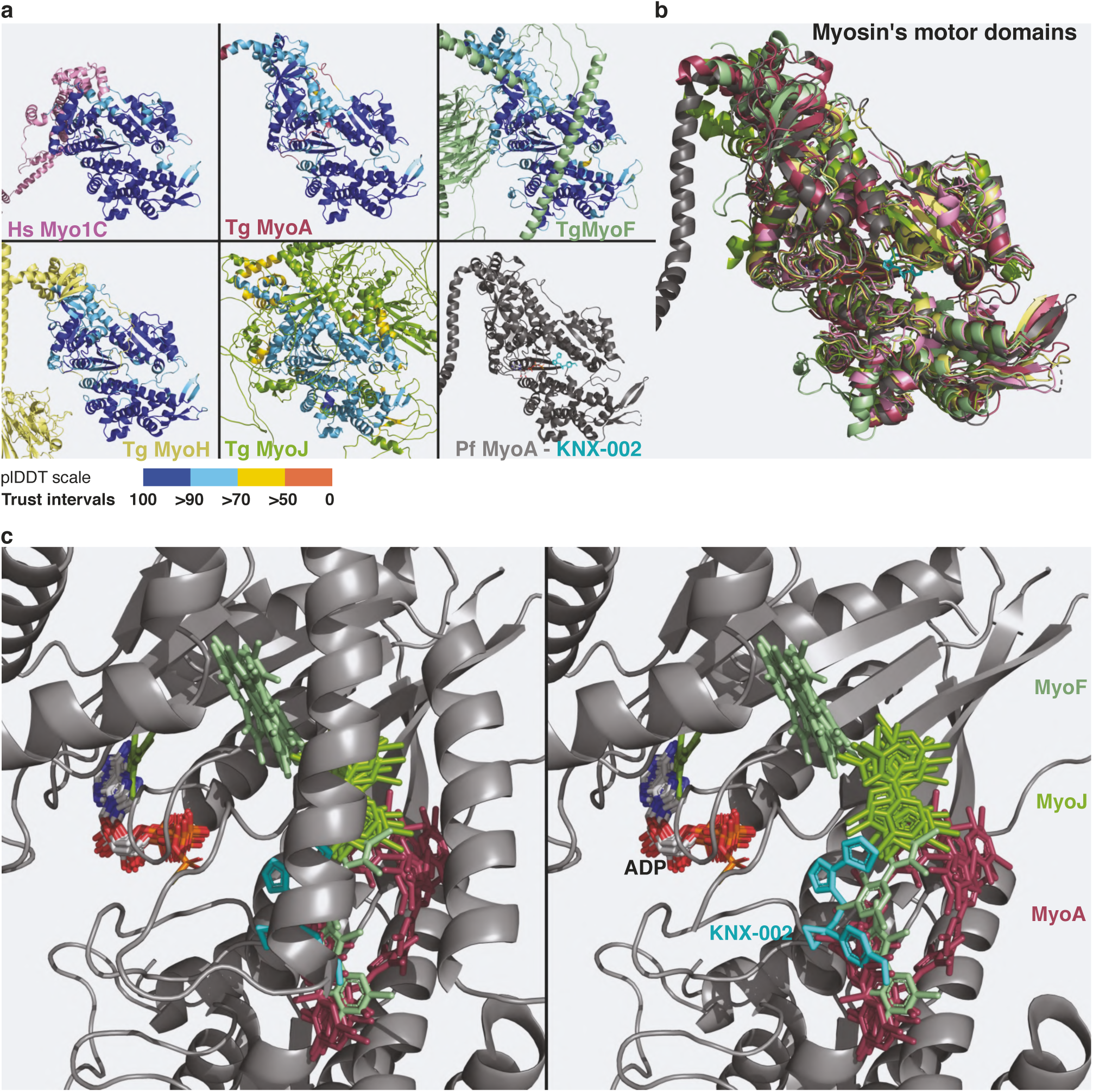
Comparison between the motor domains of human myo1C, *P. falciparum* MyoA, and several *T. gondii* myosin motor domains. **(a)** Side-by-side comparison of AlphaFold (AF3)-predicted structures of human Myo1C and *T. gondii* MyoA, MyoF, MyoH, and MyoJ, together with the crystal structure of P. falciparum MyoA (gray) bound to the MyoA inhibitor KNX-002 (cyan) and ADP (white). For each AlphaFold-predicted structure, the motor domain was used for structural alignment and is colored according to the standard pLDDT confidence scale (dark blue, very high; light blue, high; yellow, low; orange, very low). Regions outside the motor domain retain the protein-specific colors indicated above **(b)** Superposition of the motor domain structures shown in (a), using the same protein-specific color code. **(c)** Superposition of PCIP-binding poses predicted by AF3 co-folding, with or without ADP, for *T. gondii* MyoA, MyoF, MyoH, and MyoJ, aligned to the motor domain of *T. gondii* MyoF. PCIP molecules are colored according to their respective myosin partners, using the same color code as in (a). ADP molecules from the AF3 co-folding predictions are shown in white. For comparison, the experimentally resolved binding pose of the MyoA inhibitor KNX-002 (cyan) in *P. falciparum* MyoA is also shown (<u>Moussaoui et al., 2023</u>).

Having established this structural conservation of myosin motor-domain and the location of the KNX-binding site, we next investigated where PCIP might associate with *T. gondii* myosins by generating AF3 models of the MyoA, MyoF, MyoH, and MyoJ motor domains with PCIP, both in the presence and absence of ADP. When included in the modelling, ADP was consistently positioned within the expected nucleotide-binding site, at a spatially distinct site from PCIP. Importantly, models generated with PCIP alone or together with ADP populated similar PCIP pose clusters, indicating that the presence of ADP neither displaced PCIP nor promoted convergence toward a more defined binding location. Accordingly, across five independent AF3 iterations for each of the four myosins under each condition (with or without ADP), PCIP did not converge on a single reproducible binding pose shared across all myosins.

Nevertheless, recurrent patterns of PCIP placement emerged for individual myosins. In particular, PCIP occupied closely juxtaposed regions in MyoF and MyoJ, suggesting a similar preferred region of association in these two myosins. In MyoA, PCIP adopted a somewhat distinct placement but consistently localized in close proximity to the KNX-binding site (Fig. 1b, Supplementary Fig. S1a). In contrast, PCIP showed no consistent localization on MyoH and including for the majority of iterations outside from the motor domain. Therefore, MyoH was not considered further in the analysis (Supplementary Fig. S1b).

### PCIP impairs *T. gondii* tachyzoite motile behaviors

*T. gondii* tachyzoites move by an adhesion-dependent process termed gliding motility, first characterized by videomicroscopy decades ago (<u>Frixione et al., 1996</u>). On two-dimensional surfaces, tachyzoites exhibit two productive modes generating net displacement, circular and predominantly helical gliding, as well as non-productive movements while anchored at their posterior pole: rapid rotation (twirling) or side-to-side waving. In three-dimensional matrices mimicking the extracellular environment in vivo and ex vivo, tachyzoites exclusively undergo helical gliding, supporting its physiological relevance (<u>Leung et al., 2014</u>).

Recent studies have refined the MyoA-centered model of *T. gondii* helical gliding (<u>Vigetti et al., 2024</u>; <u>Hueschen et al., 2024</u>). The parasite first establishes an apical adhesion site, and generates MyoA-dependent traction to move forward while rotating ∼180°. After translocation of the cell body, it flips again and initiates a new cycle of apical attachment and forward movement. To determine whether the candidate MyoA inhibitor PCIP affects this process, we used real-time live-cell imaging of extracellular tachyzoites gliding on glass coverslips and compared vehicle-treated parasites with increasing PCIP concentrations to assess concentration-dependent effects on motility.

Tachyzoite typical helical or circular gliding was interrupted within minutes post addition of PCIP at the concentrations tested from 0.1 μM to 1 μM (Supplemental Movie 1, Fig.2a). Quantitative and statistical analysis of specific behaviors on more than 750 zoites across 3 independent assays indicate that more than 90% of the productive motile skills are lost in concentrations as low as of 0.1 μM PCIP, therefore showing high potency when compared to the KNX-115 which inhibits *P. falciparum* sporozoite gliding with an EC at about 0.117 μM (<u>Trivedi et al., 2026</u>). However, gliding was fully restored following PCIP wash out which argues for a strong but reversible target engagement and against damage of the motility machinery (Supplemental Movie 2, Fig2b).

While this phenotypic effect on motility is consistent with PCIP targeting the core motor MyoA component of the glideosome, PCIP was unlikely to completely inhibit the entire pool of MyoA activity. Therefore, we next asked whether increasing microneme secretion could compensate for the reduced motor capacity. Microneme secretion increases the availability of surface adhesins that may then engage extracellular ligands and couple to the actomyosin machinery. We therefore used the Ca²⁺ ionophore A23187 to strongly stimulate microneme secretion, reasoning that increased adhesive engagement might allow the residual myosin activity remaining under PCIP treatment to sustain motility. Remarkably, addition of A23187 to PCIP-arrested tachyzoites, while maintaining PCIP throughout the stimulation, restored motility to levels comparable to ionophore-treated controls. These findings suggest that boosting Ca²⁺-dependent microneme secretion can overcome the PCIP-induced motility defect when sufficient residual motor activity remains (Supplemental Movie 3, Fig 2c).

**Figure 2.**
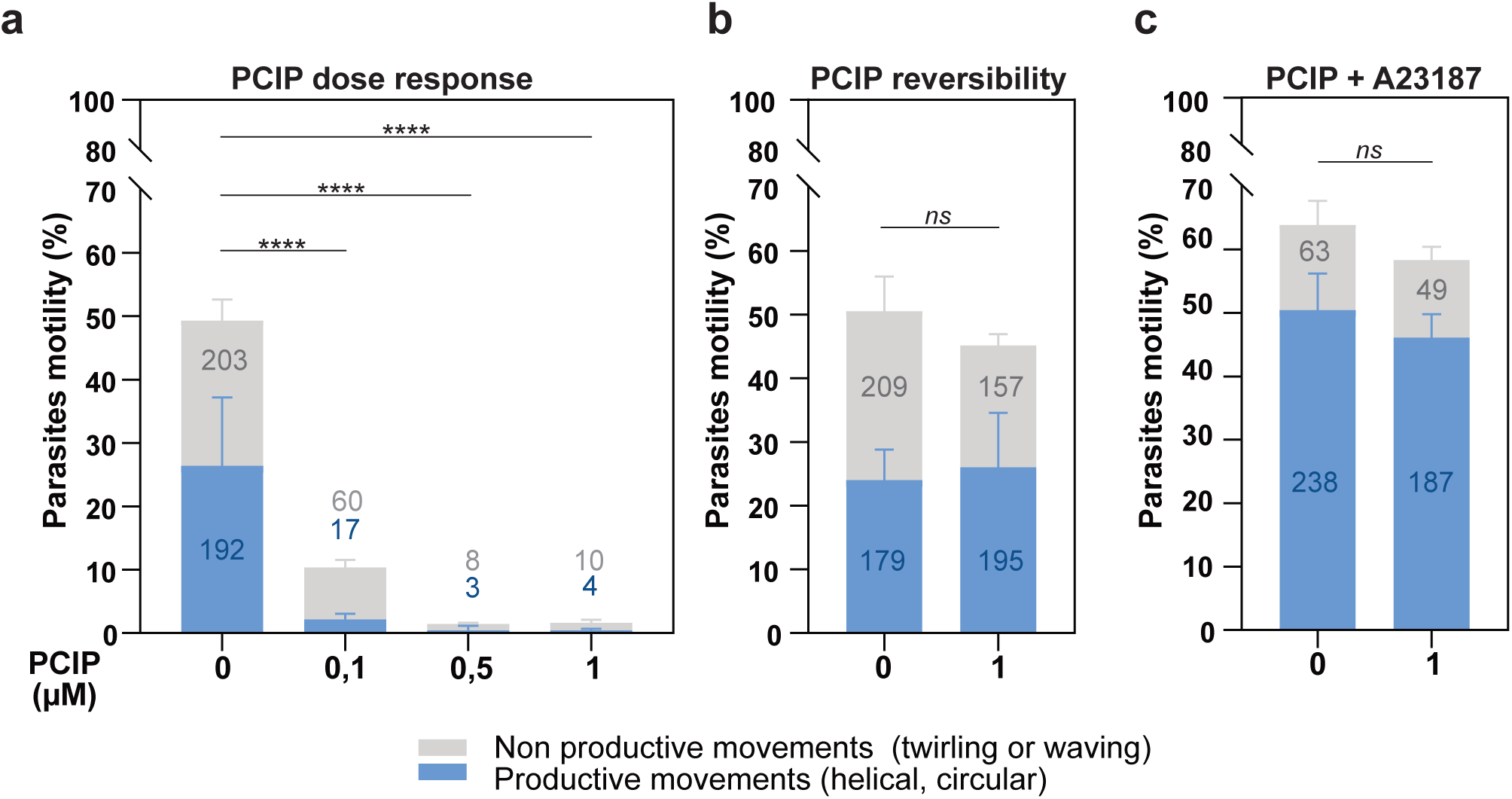
Dose-dependent and reversible inhibition of tachyzoite motility by PCIP. **(a)** Inhibition of tachyzoite productive gliding by PCIP. Motility was assessed during 5.5 min following treatment with PCIP at the indicated concentrations. The percentage of motile parasites (productive and non-productive) was quantified from n > 750 parasites across three independent assays **(b)** Recovery of tachyzoite motility following PCIP removal. Parasites were treated with PCIP and subsequently washed to remove the compound before reassessment of productive gliding. Motility was quantified after 30 min of recovery during 5.5 min. n > 750 parasites across three independent assays **(c)** Calcium ionophore A23187 counteracts the inhibitory effect of PCIP on tachyzoite motility. Parasites were first treated with PCIP to inhibit productive gliding, and the inhibitory effect of PCIP was confirmed prior to addition of the calcium ionophore A23187. Productive gliding was assessed following A23187 addition during 6 min. n > 400 parasites across two independent assays. These data indicate that an A23187-induced increase in cytosolic calcium can overcome the inhibitory effect of PCIP on tachyzoite motility. Data are presented as mean ± SD. Statistical significance was assessed using a one-way ANOVA with a significance threshold of α = 0.05. ****p < 0.0001.

### PCIP impairs *T. gondii* tachyzoite invasive behaviors

Having established that PCIP strongly impairs tachyzoite gliding, we next investigated whether inhibition of parasite motility also affected host-cell invasion, a process that relies on the acto-MyoA machinery for active penetration. Tachyzoites were briefly pre-exposed to increasing concentrations of PCIP and subsequently added to HFF monolayers in the continued presence of the corresponding PCIP concentration. Invasion was quantified by differential staining combined with AI-assisted custom image analysis (see Methods). Because tachyzoites constitutively expressed a green fluorescent marker, the total parasite population could be detected, whereas extracellular parasites were selectively labeled by SAG1 immunostaining under conditions of minimal fixative-induced cell permeabilization.

Within each experiment, the number of HFFs per field, assessed by nuclear counts, remained consistent across treatment conditions, supporting the robustness of the automated image-analysis pipeline. Macro accuracy was comparable across all three experiments, with error rates below 5% for cell and green parasite detection and slightly higher, but below 6%, for double-positive parasite detection (Supplemental Fig. S3). In contrast, the proportion of intracellular parasites progressively decreased with increasing PCIP concentrations. Repeated-measures one-way ANOVA revealed a significant overall effect of PCIP treatment (*p* = 0.0339) as detailed for each assay (Fig. S3b-d), while analysis for linear trend showed a highly significant dose-dependent decrease in host-cell invasion (*p* < 0.0001) (Fig. 3a). Consistent IC₅₀ values of approximately 0.2–0.3 µM were obtained using two independent image-analysis approaches, either a customized image-processing macro or the ScanR high-content screening platform (Supplemental Fig. 3b).

**Figure 3.**
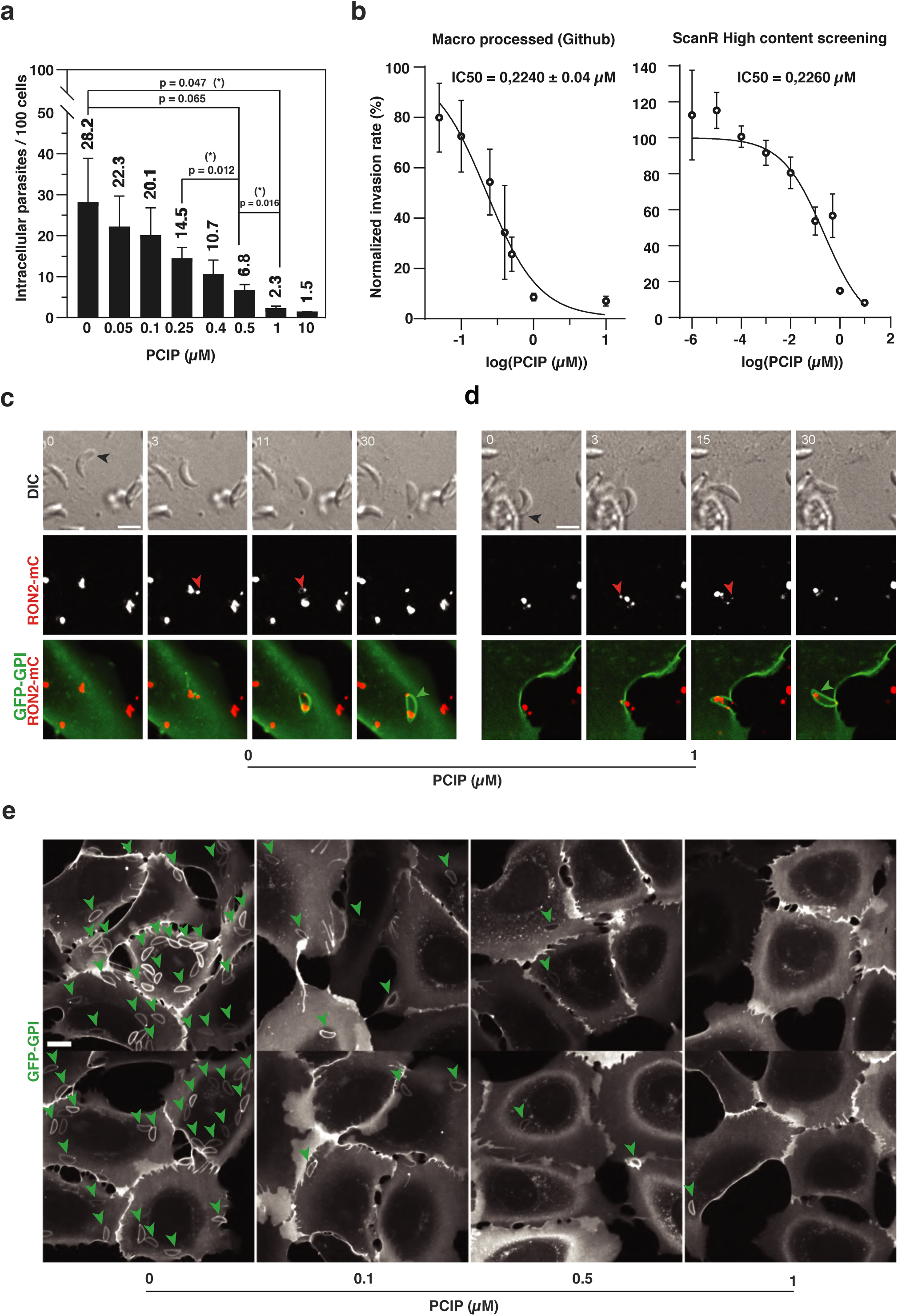
PCIP inhibits MyoA and impairs parasite invasion while preserving RON-dependent entry. **(a)** Invasion of nLuc-GFP parasites into HFF cells, expressed as the number of intracellular parasites per 100 cells, across a range of PCIP concentrations in µM in three independent assays. Values were quantified from > 30,000 cells in the control condition, around 15,000 cells per PCIP condition (0.05-10 µM), and around 10,000 cells for 0,4 and 10 µM. Bars show the mean ± SD of n = 3 assays. p-values are indicated. Mixed-effects model’s linear trend was highly significant (p-value < 0,0001) of a PCIP dose-dependent reduction in invasion. **(b) Left** : Dose-response curves and IC50 estimation across two different independent experimental conditions and image-processing techniques: IC50 estimation in the three invasion assays processed with the customized Fiji macro available on GitHub. **Right**: an additional assay with broader concentration ranges, analysed by ScanR high-content screening. The two pipelines gave near-identical IC_50_ values (0.224 µM vs 0.226 µM) **(c-d)** Time-lapse imaging of DIC (top) and fluorescent channel overlays (bottom) showing a RON2-mC tachyzoite (red) invading a U2OS cell expressing a GFP-GPI PM marker (green) under vehicle **(c)** or PCIP (1 µM) conditions **(d)**. The RON2-mC invasive nanodevice is injected into the host-cell PM, where it assembles into a torus (red arrows) through which the parasite enters in a budding PV is indicated with green arrowheads. Invasion at 1 µM PCIP occurs through similar kinetics and features than untreated parasites. N = 2 independent assays. Time is indicated in seconds. Scale bar, 5 µm. **(e)** Representative images acquired after 17 min of live imaging of RON2-mC tachyzoites (red) invading U2OS cells expressing a GFP-GPI plasma membrane (PM) marker (green), under vehicle conditions or in the presence of PCIP (0.1, 0.5, and 1 µM). Green arrows indicate PVs. N = 2 independent assays. Scale bar, 10 µm.

We previously showed that, in the absence of MyoA, tachyzoites can nevertheless enter host cells and establish a replication-permissive parasitophorous vacuole (PV) through an alternative invasion mechanism in which host-cell cortical actin contributes to the force required for parasite internalization (<u>Bichet et al., 2016</u>). We therefore investigated whether the residual invasion observed in the presence of PCIP involved a similar alternative mode of entry. Live-cell imaging was used to examine the dynamics of individual invasion events under 1 μM PCIP expected to impair MyoA-dependent motility.

Consistent with the quantitative invasion assay, successful invasion events were markedly less frequent in the presence of PCIP than under control conditions. However, the residual invasion events captured by live imaging retained features characteristic of the conventional active mode of entry (Fig. 3c, Supplemental Movie 4). Parasites expressing the chimeric rhoptry neck protein RON2-mCherry (RON2-mC) released the RON2 complex at the target cell membrane and squeezed through the host–parasite junction while entering a nascent PV, whose membrane was visualized using the host-cell GFP-GPI membrane marker. Representative images acquired at the end of the 17-min recording sequences show parasites enclosed within GFP-positive parasitophorous vacuole membranes (PVMs) and illustrate the significant reduction in invasion events with increasing PCIP concentrations (Fig. 3d). Although PCIP markedly reduced invasion efficiency, the few invasion events that persisted showed no obvious switch to the previously described host-driven internalization mode. Instead, their dynamics remained consistent with residual acto-MyoA-dependent entry, suggesting that sufficient motor activity may persist at or near the cell-tachyzoite junction to support invasion in a minimal subset of PCIP-treated parasites.

Together with the motility phenotype, these results indicate that pharmacological inhibition of the tachyzoite gliding motor by PCIP strongly compromises both gliding motility and host-cell invasion.

### PCIP inhibits intracellular replication of *T. gondii* tachyzoites at submicromolar concentrations without affecting host cell viability

Tachyzoites replicate by endodyogeny, a process in which two daughter cells are assembled within the mother parasite with their apicobasal axes most frequently aligned toward the maternal apex (<u>Hu et al., 2002</u>). Recent work has shown that apico-basal polarity is established very early during endodyogeny, concomitantly with daughter bud formation (<u>Padilla et al., 2024</u>). MyoF contributes to this process by ensuring proper centrosome positioning following duplication, thereby defining the site of daughter bud emergence (<u>Jacot et al., 2013</u>). In addition, MyoF performs several other essential functions during early endodyogeny, including apicoplast segregation (<u>Devarakonda et al., 2023</u>) and the positioning of multiple endomembrane compartments such as the apical secretory organelles. We therefore investigated whether these MyoF-dependent processes during intracellular tachyzoite development are affected by PCIP.

To discriminate between effects on host-cell invasion and intracellular replication, GFP-expressing *T. gondii* tachyzoites were first allowed to invade pseudo-confluent or confluent A549 or HFF monolayers. PCIP was then added at increasing concentrations 15 min after completion of invasion. Intracellular parasite development was assessed 22 h post-infection by quantifying the number of tachyzoites per PV using GFP fluorescence together with Hoechst DNA staining (Fig. 4a). In untreated cultures, most PVs contained eight parasites, with a proportion already reaching sixteen parasites, indicating that tachyzoites had completed three to four rounds of endodyogeny during the 22 h incubation period. A similar distribution was observed following treatment with 0.1 µM PCIP, indicating that intracellular replication was unaffected within this time period. In contrast, increasing PCIP concentrations progressively impaired parasite division. At 0.4 (data not shown) and 0.5 µM, most PVs contained either a single parasite, or two daughter parasites, suggesting that replication was arrested during or early after the first round of endodyogeny. At 1 µM PCIP, nearly all parasites failed to complete their first division cycle (Fig. 4a). Quantitative analysis confirmed that PCIP treatment significantly altered the distribution of parasites per vacuole (p = 0.0076) and specifically between parasites exposed to vehicle and 0.5 µM PCIP (p = 0.0013), to 0.1 and 0.5 µM (p = 0.0141), and to 0.5 and 1 µM (p = 0.0174).

**Figure 4.**
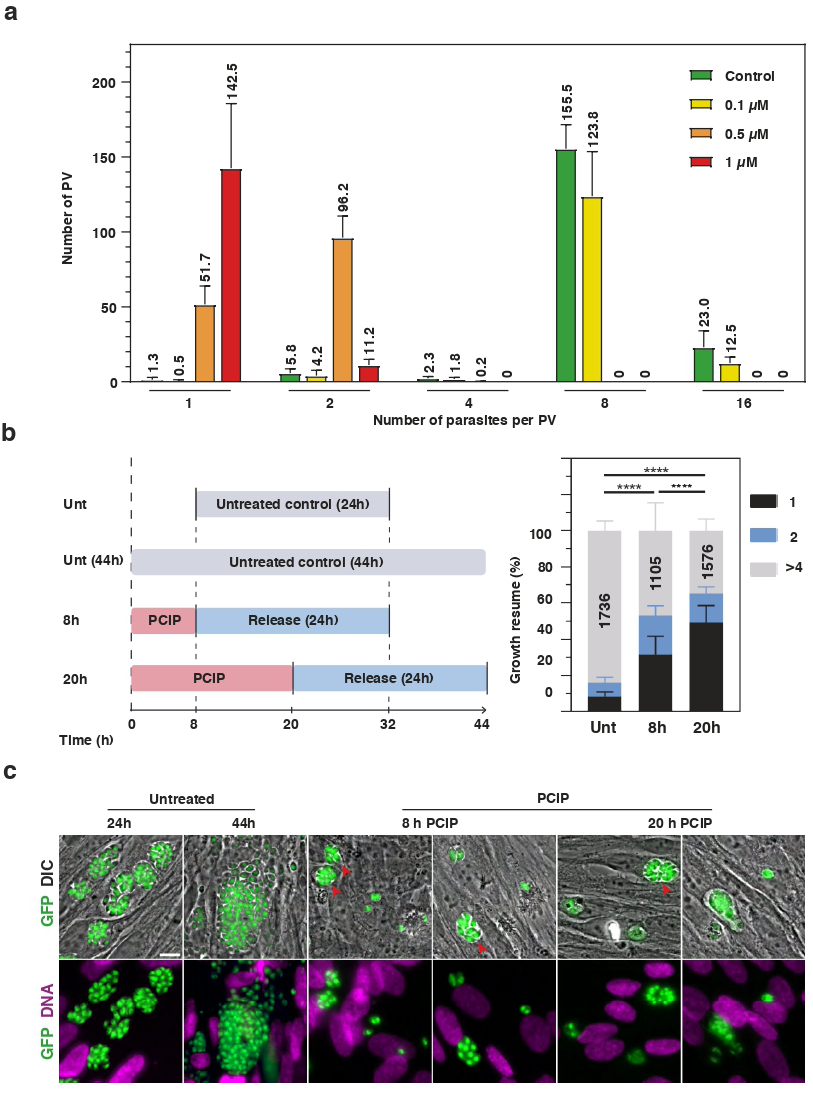
PCIP inhibits intracellular replication of *T. gondii* tachyzoites in a partially reversible manner. **(a)** Histogram showing the distribution of the number of tachyzoites (1, 2, 4, 8, or 16) per PV at 22 h post-invasion. Following a 15-min invasion period, extracellular parasites were removed and infected cells were exposed to vehicle (DMSO; untreated, green) or PCIP at 0.1 µM (yellow), 0.5 µM (orange), or 1 µM (red). A minimum of 250 PVs per replicate were analyzed for each condition in three independent experiments. Analysis of parasite distribution per PV across all conditions revealed a significant treatment × parasite-number category interaction (p = 0.0076). The distribution of parasites per PV was significantly shifted toward lower parasite numbers, consistent with impaired endodyogeny, in parasites exposed to 0.5 µM PCIP compared with vehicle-treated parasites (p = 0.0013) or parasites exposed to 0.1 µM PCIP (p = 0.0141). Replication was further impaired at 1 µM PCIP compared with 0.5 µM PCIP (p = 0.0174). **(b)** Assessment of parasite growth recovery following PCIP removal. **Left:** Schematic representation of the washout protocol. Intracellular parasites were exposed to 0.5 µM PCIP for 8 or 20 h, after which PCIP was removed and parasites were cultured in drug-free medium for an additional 24 h. Vehicle-treated (DMSO) controls were included at t = 0 and at the corresponding washout time points to compare parasite growth during the subsequent 24-h recovery period. **Right:** Histograms showing the distribution of parasite progeny (1, 2, or ≥4 tachyzoites) per PV after the 24-h recovery period for vehicle-treated controls (Unt) and parasites exposed to 0.5 µM PCIP for 8 or 20 h. Chi2 test showed that transient PCIP treatment highly significantly decreased the amount of progeny per vacuole in a time dependent way (p < 0.0001). **(c)** Representative epifluorescence images of HFFs infected with GFP-expressing tachyzoites following the experimental conditions described in **(b)**. Images show vehicle-treated controls and parasites exposed to 0.5 µM PCIP for 8 or 20 h, followed by PCIP washout and a 24-h recovery period in drug-free medium. The top panels show overlays of transmitted-light and fluorescence images, with GFP fluorescence identifying the tachyzoites. The bottom panels show overlays of green and UV fluorescence, allowing visualization of tachyzoites and the nuclei of both parasites and host cells, respectively. Red arrowheads mark the growth-permissive PVs. Scale bar, 10 µm.

To determine whether the effects of PCIP on tachyzoite growth could result from a detrimental effect on host cells, we assessed PCIP cytotoxicity in the immortalized human A549 epithelial cell line and in HFFs, which become contact-inhibited and exhibit limited proliferation upon reaching confluence. No increase in SYTOX Green-positive cells was detected following 24 h of treatment at concentrations up to 80 μM, indicating that PCIP did not induce detectable cell death under these conditions. In contrast, quantification of Hoechst-stained nuclei revealed a concentration-dependent reduction in cell number in both cell type but slightly more in the actively proliferating A549 cells. These findings are consistent with an inhibition of cell proliferation at doses higher than 1 μM rather than cytotoxicity (SupTable 1).

### The PCIP-induced arrest of tachyzoite proliferation is only partially reversible

To determine whether PCIP exerts a parasitostatic or parasiticidal effect on intracellular *T. gondii*, we assessed the ability of arrested parasites to resume replication following drug removal. HFF cells infected with tachyzoites were treated with PCIP for 8 or 20 h at a concentration shown to inhibit endodyogeny (i.e., 0.5 mM) and subsequently cultured in drug-free medium for an additional 24 h (Fig. 4b). Following washout after 8 or 20 h of treatment, approximately 50% and 35% of the arrested parasites resumed intracellular replication, respectively (t = 0, at which point egress had already begun; t = 8 h; and t = 20 h) (Fig. 4b). In contrast, parasites that failed to recover mostly remained as single or paired intracellular tachyzoites, often displaying only a faint GFP signal, consistent with a loss of viability (Fig. 4c).

The number of replicating tachyzoites within PVs was significantly reduced following 8 h of PCIP treatment and subsequent washout (p < 0.0001). When PCIP treatment was prolonged to 20 h, growth recovery was significantly lower than that observed after the shorter treatment (p < 0.0001) (Fig. 4b, graph). Together, these results indicate that PCIP-induced arrest of endodyogeny is only partially reversible, suggesting that PCIP exerts both parasitostatic and parasiticidal effects on intracellular parasites.

### PCIP blocks early daughter cell scaffolding during live imaging of endodyogeny

To characterize PCIP effects on endodyogeny with high temporal and spatial resolution, we performed live-cell imaging in infected A549 and HFF cells. Daughter-cell formation was monitored using IMC12-mCherry (IMCmC), which labels the inner membrane complex (IMC) outlining developing daughters (<u>Vigetti et al., 2022</u>), and BCC7-mEmerald (BCC7Em) (<u>Engelberg et al., 2016</u>). BCC7 is recruited to daughter bud initiation sites, then moves along the growing daughter periphery before accumulating as a ring at the posterior pole in association with Myo C (<u>Vigetti et al., 2022</u>).

Consistent with quantitative growth assays, ∼30-h live imaging showed similar daughter-cell formation and rosette organization in vehicle- and 0.1 µM PCIP-treated parasites. At 0.1 µM, IMC12 and BCC7 dynamics during daughter assembly resembled controls (Fig. 5a,b; Supplemental Movie 5). Maternal IMC12 progressively disassembled toward the basal pole, while cytosolic IMC12 reorganized into the two daughter IMC scaffolds. BCC7 appeared as apical foci in nascent daughters, moved basally as the IMCs elongated, and accumulated at daughter basal poles. In contrast, at 0.5 µM PCIP, most tachyzoites remained arrested before endodyogeny throughout the 21-h recording, with neither IMC12 nor BCC7 showing these dynamic redistributions (Fig. 5c; Supplemental Movie 6).

**Figure 5.**
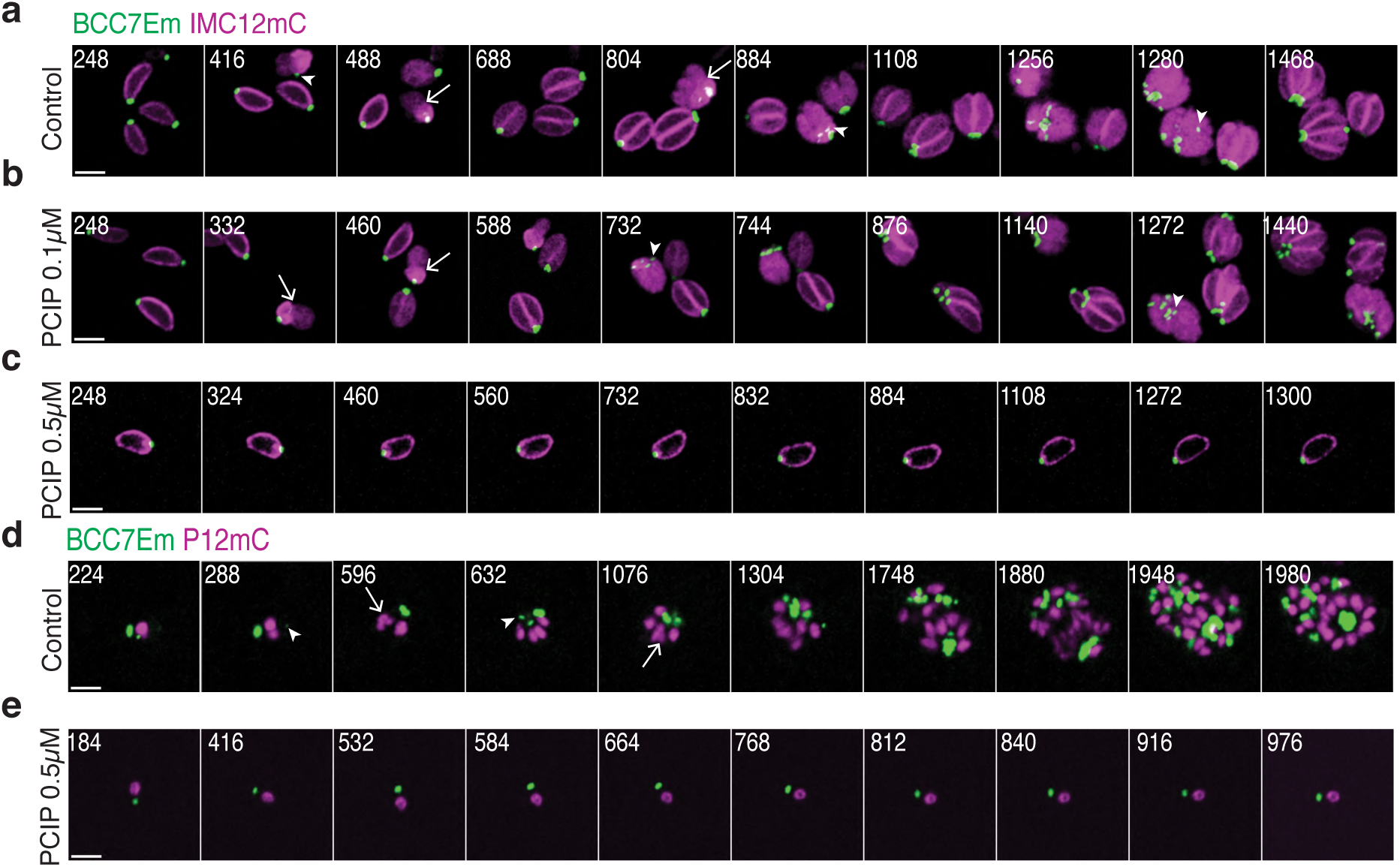
PCIP blocks early daughter-cell scaffolding during live-cell imaging of endodyogeny. Selected frames from live cell time-lapse imaging of intracellular *T. gondii* tachyzoites expressing fluorescent reporters of daughter-cell formation, in the absence or presence of PCIP. The time corre-sponding to each selected frame is indicated in min **(a)** Untreated control parasites undergo typical en-dodyogeny, illustrated by the dynamic redistribution of IMC12-mCherry, which progressively disassembles from the maternal cytoskeleton and collapses toward the posterior residual body (white open arrows). Concomitantly, Basal Complex Component 7 (BCC7) undergoes anteroposterior redistribution (white arrowheads), while cytoplasmic IMC components assemble into the characteristic IMC-enriched scaffolds of the two emerging daughter cells **(b)** Similar dynamic changes in IMC12 and BCC7 localization are observed in intracellular tachyzoites exposed to 0.1 µM PCIP, indicating that endodyogeny remains active at this concentration **(c)** In contrast, no detectable dynamics of IMC12 or BCC7 are observed during 30 h of live-cell imaging of tachyzoites exposed to 0.5 µM PCIP, consistent with the absence of daughter-cell scaffold formation **(d)** Untreated control para-sites expressing the endogenous nuclear marker P12-mCherry undergo the characteristic change in nuclear morphology from a spherical to a U-shaped configuration (white open arrows) that accompanies BCC7 trafficking (white arrowheads) and early daughter-cell budding **(e)** In contrast, intracellular parasites exposed to 0.5 µM PCIP fail to divide and typically retain a rounded nucleus, with no detectable transition to the U-shaped nuclear morphology associated with daughter-cell formation.

To further determine whether PCIP arrests parasites before or after daughter bud initiation, we monitored nuclear morphology as a readout of endodyogeny progression. Following centrosome duplication, the nucleus remains rounded during bud initiation and becomes characteristically U-shaped as daughter scaffolds elongate. Nuclear dynamics were therefore co-monitored with daughter assembly using a mCherry fusion of a 15.9-kDa nuclear matrix protein previously identified as partner 12 in the BCC7 interactome (<u>Vigetti et al., 2022</u>). The U-shaped nucleus preceding nuclear fission was readily observed during successive divisions in vehicle-treated parasites (Fig. 5d; Supplemental Movie 7), but not at 0.5 µM PCIP, even during recordings exceeding 20 h (Fig. 5e; Supplemental Movie 8).

### Super-resolution microscopy characterizes the PCIP-induced block in tachyzoite replication

Ultrastructure expansion microscopy (U-ExM) has proven particularly effective for resolving the inner membrane complex (IMC) and subpellicular microtubules of *T. gondii* (<u>Pavlou et al., 2020</u>; <u>Tosetti et al., 2020</u>; <u>Padilla et al., 2024</u>). We therefore used U-ExM to examine the structural organization of tachyzoites following PCIP treatment. Infected-cell samples were processed for U-ExM and subjected to post-expansion staining with previously validated antibodies against either *T. gondii* IMC1 or acetylated α-tubulin, while nuclear DNA was visualized using propidium iodide.

At 24 h post-infection, cells treated with either DMSO or 0.1 µM PCIP contained vacuoles harboring 8-16 tachyzoites (Fig. 6, Fig. 7). Mature parasites arranged in rosettes were readily identified by their continuous peripheral IMC (Fig. 6a,b; top panel). Parasites undergoing endodyogeny were also observed, with two nascent daughter IMC scaffolds developing within the maternal IMC (Fig. 6ab; bottom panel). The characteristic U-shaped morphology of the dividing nucleus provided an additional hallmark of coordinated nuclear division and daughter-cell budding (Fig 6a,b; bottom panels).

**Figure 6.**
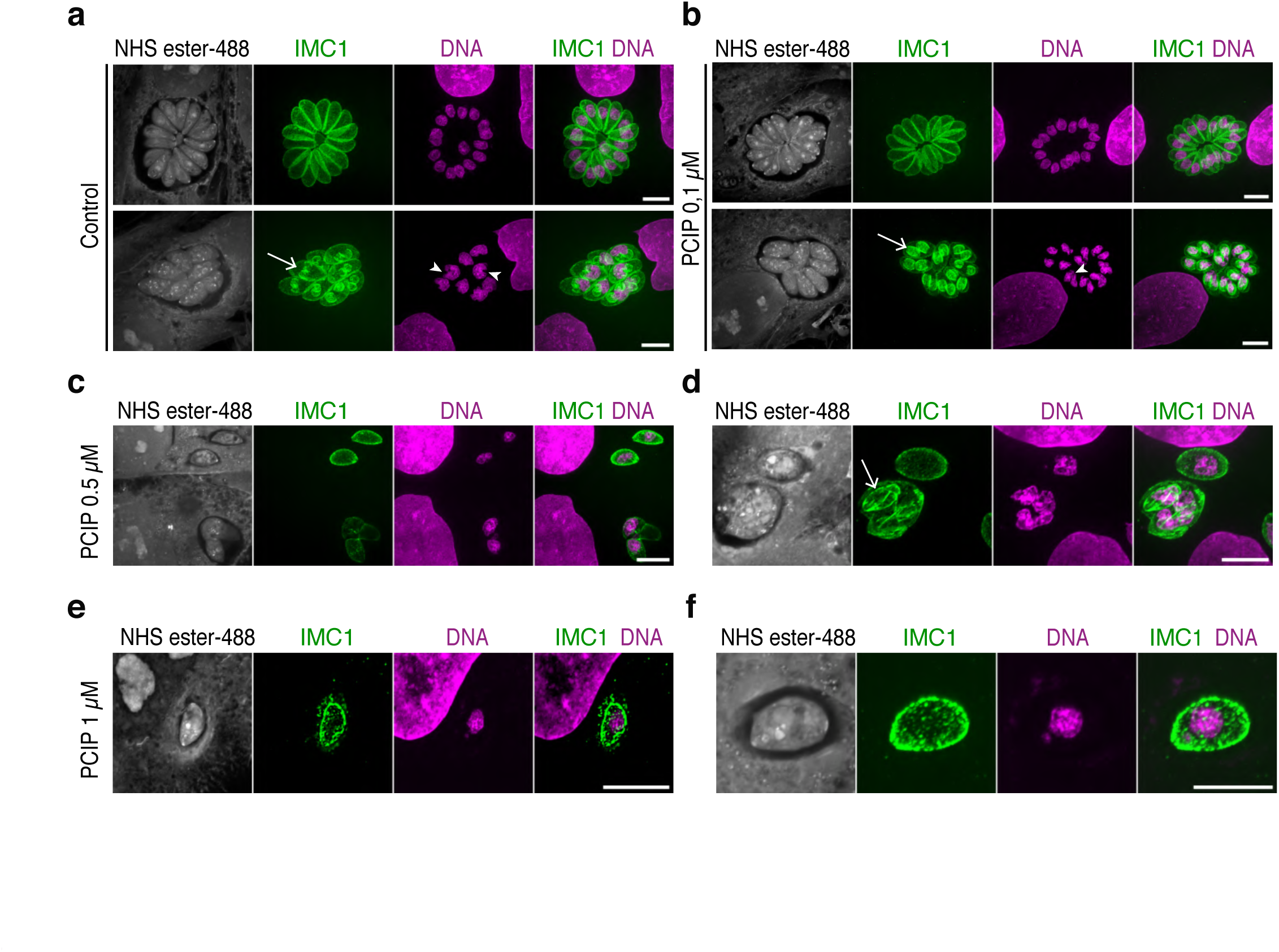
Super-resolution microscopy reveals a PCIP-induced block in IMC dynamics and daughter-cell formation during tachyzoite endodyogeny. HFFs infected with *T. gondii* tachyzoites were exposed for 24 h to vehicle control **(a)** or increasing concentrations of PCIP **(b-f)** and analyzed by ultrastructure expansion microscopy (U-ExM). NHS-ester staining reveals the overall cellular organization, while IMC1 staining highlights the peripheral inner membrane complex (IMC) of mature parasites and the developing IMC scaffolds of daughter cells **(a, b).** In vehicle-treated parasites **(a)** and parasites exposed to 0.1 µM PCIP **(b)**, the top panels show mature tachyzoites organized in typical rosettes and displaying peripheral IMC1 staining. The corresponding bottom panels show parasites undergoing endodyogeny, with IMC1 delineating the scaffolds of emerging daughter cells within the mother cell (white open arrows). Daughter-cell formation is accompanied by the characteristic fork-shaped nuclear morphology (white arrowheads) associated with nuclear material entering each nascent daughter bud. **(c, d)** At intermediate PCIP concentrations, most parasites fail to divide. However, occasional parasites display aberrant, apparently abortive budding, characterized by misshapen IMC-positive daughter scaffolds and abnormal organization of the nuclear material. **(e, f)** At the highest PCIP concentrations, including 1 µM, parasites are predominantly found as single tachyzoites retaining peripheral IMC1 staining and a largely spherical nucleus, with no evidence of daughter-cell scaffold formation. These phenotypes are consistent with a severe impairment of endodyogeny. Scale bars, 5 µm.

**Figure 7.**
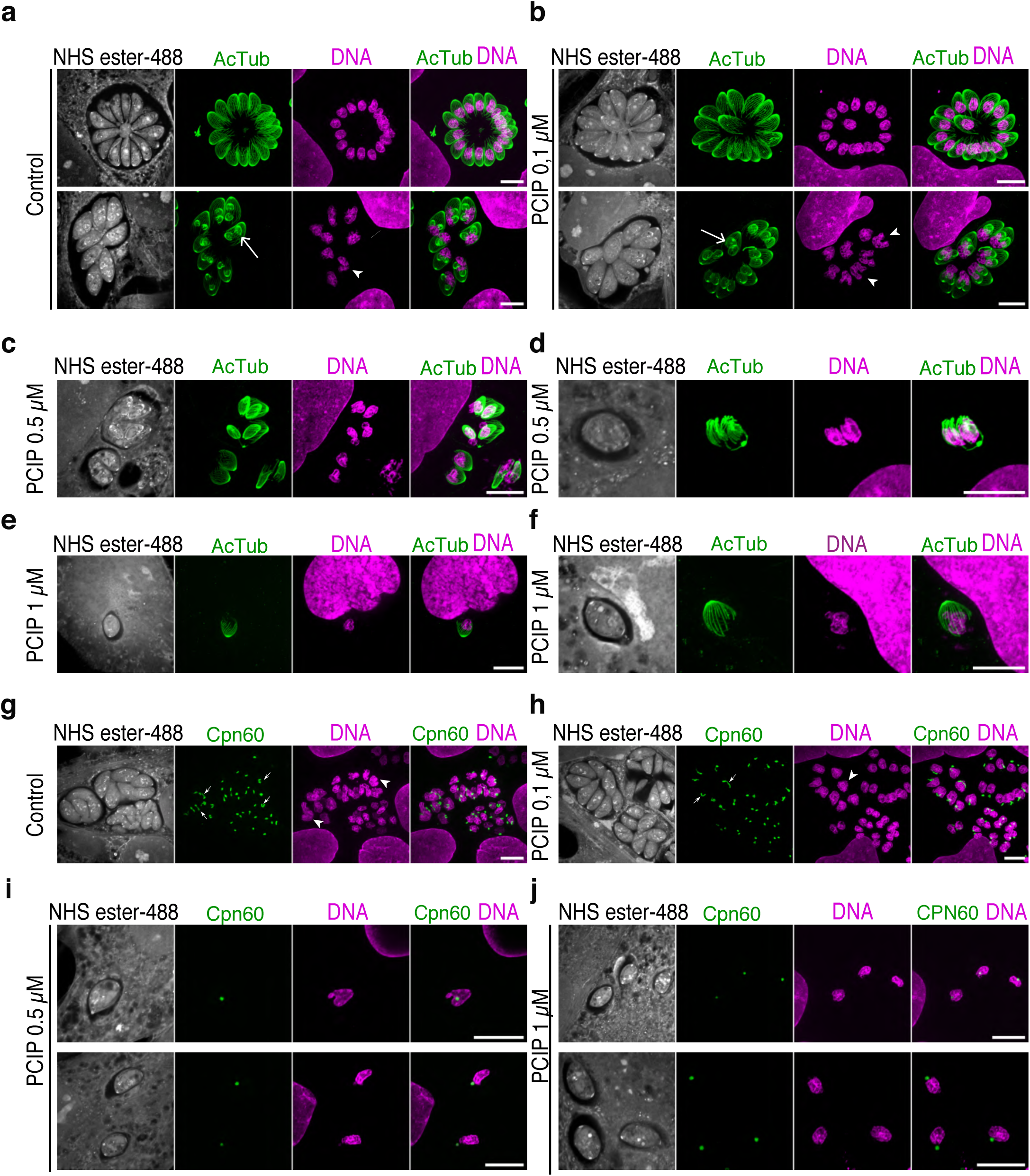
Super-resolution microscopy reveals PCIP-induced defects in cortical microtubule assembly and apicoplast division during tachyzoite endodyogeny. HFFs infected with *T. gondii* tachyzoites were exposed for 24 h to vehicle control **(a, g)** or increasing concentrations of PCIP **(b–f, h–j)** and analyzed by ultrastructure expansion microscopy (U-ExM). NHS-ester staining reveals the overall cellular organization **(a–j)**, while acetylated-tubulin and apicoplast staining were used to monitor cortical microtubule (cMT) organization **(a–f)** and apicoplast dynamics **(g–j)**, respectively.**(a, b)** In vehicle-treated parasites **(a)** and parasites exposed to 0.1 µM PCIP **(b)**, acetylated-tubulin staining highlights the cortical microtubule network and a prominent ring-like structure at the parasite apex, consistent with the apical polar ring (APR) from which the 22 cMTs originate. Mature parasites display the characteristic peripheral cMT basket (top panels). In parasites undergoing endodyogeny (bottom panels), acetylated-tubulin staining reveals the early apical rings of the nascent daughters and the cMT scaffolds extending from these structures (white open arrows). Daughter-cell formation is accompanied by the characteristic fork-shaped nuclear morphology (white arrowheads), consistent with ongoing endodyogeny. **(c, d)** At intermediate PCIP concentrations, occasional parasites initiate daughter-cell formation but display aberrant organization of acetylated-tubulin-positive apical structures and cMTs within the developing buds, resulting in misshapen or apparently abortive daughter scaffolds. **(e, f)** At higher PCIP concentrations, parasites predominantly remain as single mother cells and retain their pre-existing peripheral cMT basket and apical acetylated-tubulin-positive ring, with no detectable assembly of daughter cMT scaffolds, consistent with a block in the initiation or progression of endodyogeny. **(g-j)** Apicoplast dynamics during endodyogeny in vehicle- and PCIP-treated parasites. In vehicle-treated parasites **(g)** and parasites exposed to 0.1 µM PCIP **(h)**, the apicoplast undergoes the characteristic morphological changes associated with early endodyogeny: the organelle elongates and extends between the developing daughter cells before undergoing fission, allowing segregation of one apicoplast into each daughter (white arrows). At higher PCIP concentrations **(i, j)**, the apicoplast fails to undergo this characteristic elongation and division and instead remains as a single, undivided organelle. The nucleus likewise remains single and displays either a spherical or elongated morphology, with no successful progression to daughter-cell formation. Together, these observations indicate that PCIP disrupts both daughter cMT scaffold assembly and apicoplast division during endodyogeny. Scale bars, 5 µm.

By contrast, organized rosettes of proliferating tachyzoites were never observed following treatment with 0.5 or 1 µM PCIP (Fig. 6c-f). Instead, PV contained single parasites or parasite pairs, in which limited IMC buds were occasionally observed. Consistent with these findings, mature parasites exposed to vehicle or 0.1 μM of PCIP displayed a typical basket of 22 subpellicular microtubules (Fig. 7a,b; top panels) and emerging arrays of subpellicular microtubules were detected at the site of daughter-cell budding, indicative of ongoing endodyogeny (Fig. 7a,b; bottom panels). In contrast, disorganized, and likely abortive MT buds were observed in samples treated with 0.5 μM of PCIP (Fig. 7c,d), whereas no daughter-associated MT buds were detected in the U-ExM images acquired from samples treated with 1 μM PCIP (Fig. 7e,f). At this higher concentration, tachyzoites appeared swollen within their PV. These observations are consistent with a PCIP-induced block occurring before, or at the onset of, daughter-cell budding.

Successful apicoplast division and inheritance depend MyoF, which regulates the elongation dynamics required for the organelle to associate with duplicated centrosomes and thereby ensures that each daughter cell receives an apicoplast (<u>Devarakonda et al., 2023</u>). Samples processed for were post-stained using antibodies against the luminal chaperonin Cpn60. Dumbbell-shaped elongated apicoplasts or paired apicoplasts associated with U-shaped dividing nuclei were observed in parasites treated with vehicle or 0.1 µM PCIP (Fig. 7g,h) consistent with ongoing endodyogeny. By contrast, parasites exposed to higher PCIP concentrations (0.5 and 1 µM) contained predominantly single, rounded apicoplasts (Fig. 7i,j), providing further evidence that PCIP interferes with an early stage of the division process.

### PCIP compromises the late step of progeny maturation prior to egress

During the final phase of endodyogeny, the daughter IMCs extend toward the posterior pole and the basal complexes constrict, producing the tapered shape of mature tachyzoites. This localized contractile process involves TgMyoJ and the recruitment of TgCentrin2 (<u>Hu, 2008</u>; <u>Frenal et al., 2017</u>). Concurrently, the maternal cytoskeleton is dismantled, organelle segregation is completed, and the mature daughters remain connected through their basal poles to the residual body by cytoplasmic bridges associated with TgMyoI and F-actin, thereby maintaining the rosette organization. Before egress, the intravacuolar F-actin network collapses and the parasites detach from the residual body and become individualized and competent for MyoH–MyoA-dependent motility (<u>Periz et al., 2017</u>). Although MyoI and MyoJ are dispensable for tachyzoite growth *in vitro*, their deletion disrupts parasite connectivity, rosette organization and basal-complex maturation, respectively.

Live imaging using the IMC12-mC reporter suggested that, over longer recording periods, the growth of some parasites exposed to 0.1 µM PCIP was compromised (Supplemental Movie 5, see asterisk) and that egress appeared less efficient than in vehicle-treated parasites (data not shown). However, the extensive remodeling of the inner membrane complex (IMC) accompanying parasite replication resulted in highly dynamic changes in the IMC12-mC signal, making parasite death or lysis difficult to ascertain reliably. We therefore performed prolonged live imaging of parasites expressing cytoplasmic GFP, which provides a comparatively stable, non-dynamic fluorescence signal and thus a more reliable readout of parasite integrity and lytic events. GFP-expressing parasites were allowed to replicate intracellularly for about 30 h under culture conditions and were subsequently imaged for approximately 24 h to monitor their fate under PCIP treatment. Under control conditions, most PVs supported additional rounds of daughter-cell formation followed by coordinated egress, and invasion by the newly released tachyzoites was also observed (Fig. 8a, top panel; Supplemental Movie 9). By contrast, in a substantial proportion of PVs continuously exposed to 0.1 µM PCIP, daughter tachyzoites released their GFP-rich cytosol into the PV lumen before egress. The GFP signal remained confined within the PV, indicating a loss of parasite plasma-membrane integrity while the PV membrane remained intact (Fig. 8a, bottom panel, red arrowheads; Supplemental Movie 10).

**Figure 8.**
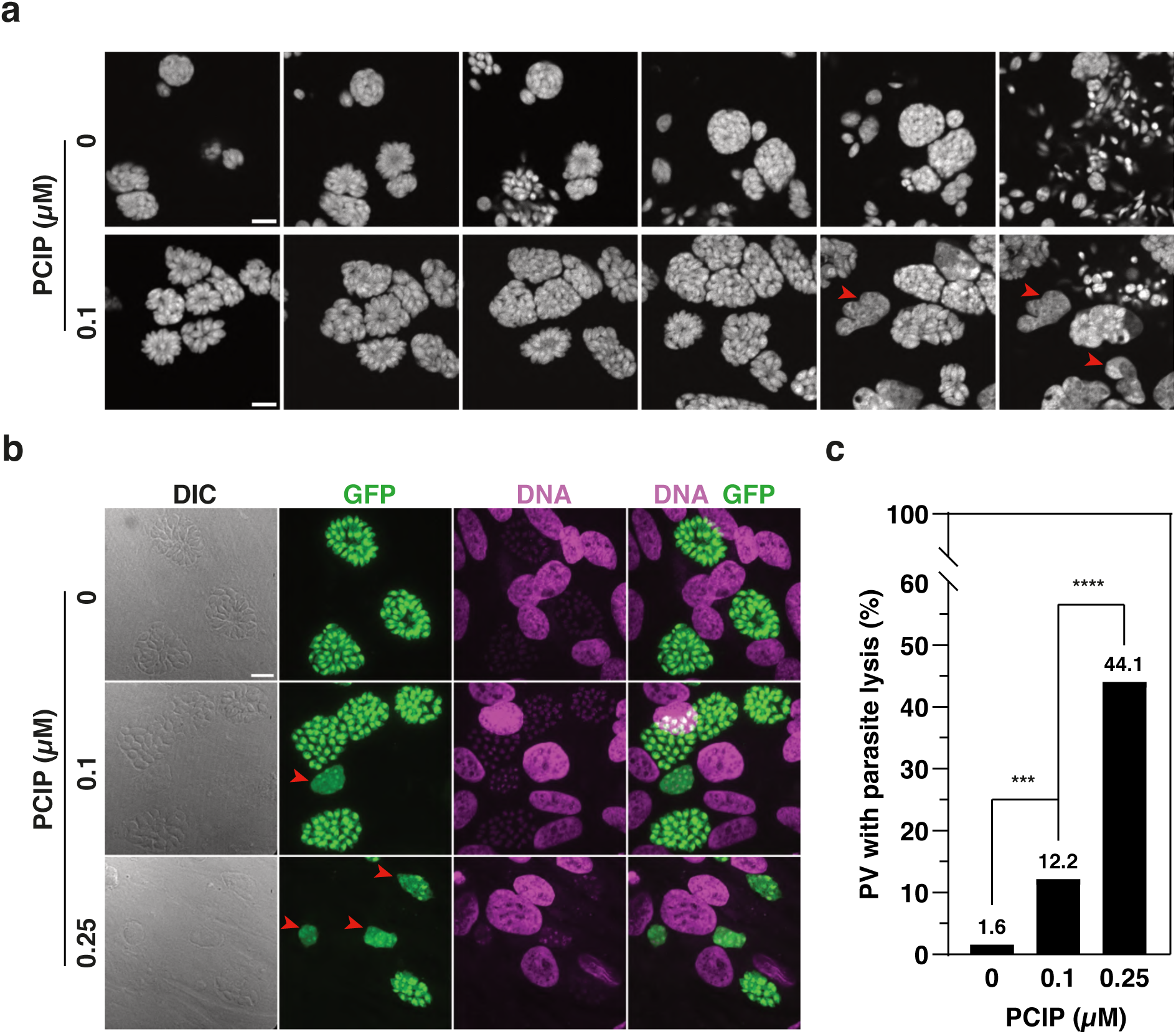
PCIP induces intravacuolar lysis (“implosion”) of *T. gondii* tachyzoites. **(a)** Selected frames from live-cell time-lapse imaging of GFP-expressing tachyzoites replicating in HFF cells treated with vehicle (0 µM PCIP; top) or 0.1 µM PCIP (bottom). The time corresponding to each selected frame is indicated in hours (h :min). Vehicle-treated parasites undergo successive rounds of endodyogeny and expand within PVs, whereas PCIP treatment induces parasite lysis at later time points, accompanied by the release and diffusion of GFP into the vacuolar space (red arrowheads). Scale bar, 10 µm. **(b)** Representative confocal images of GFP-expressing tachyzoites (green) within PVs after 30–36 h of treatment with 0, 0.1, or 0.25 µM PCIP. Host-cell and parasite nuclei were stained with Hoechst (blue). Red arrowheads indicate PVs containing lysed (“imploded”) parasites. Scale bar, 10 µm. **(c)** Proportion of PVs displaying tachyzoite implosion following treatment with 0, 0.1, or 0.25 µM PCIP. A total of 125–165 PVs per condition were analyzed across 10 different fields from a single experiment. Statistical significance was assessed using a one-sided Fisher’s exact test (\*\**p* = 0.0003; \*\*\**p* < 0.0001).

Because this phenotype emerged during prolonged recordings, we examined whether it could be exacerbated by imaging-associated phototoxicity. GFP-expressing tachyzoites were therefore cultured with PCIP for 30–36 h without live imaging. Loss of parasite-confined GFP, indicative of tachyzoite lysis, was again detected (Fig. 8b, red arrowheads), and quantified across the samples (Fig. 8c), confirming that the phenotype was not dependent on video-recording conditions. Of note, this late loss of progeny integrity was statistically more pronounced at 0.25 µM PCIP, supporting a concentration-dependent effect (Fig. 8c). Thus, at the population level, PCIP exposure reduced the proportion of PVs undergoing successful egress, consistent with an overall impairment of egress efficiency.

## Conclusions

Targeting myosin-dependent processes with small-molecule inhibitors represents a promising strategy against Apicomplexa, and *T. gondii* provides a powerful model to evaluate this approach. Its complete tachyzoite lytic cycle can be reproduced *in vitro*, allowing quantitative analysis of motility, invasion, intracellular replication and egress. Moreover, *T. gondii* expresses well-characterized unconventional myosin species with essential functions throughout this cycle.

Although inhibitors targeting MyoA-dependent motility and invasion have been identified, the pharmacological vulnerability of other myosin-dependent processes during intracellular development remains largely unexplored. Here, we dissected the effects of PCIP across the lytic cycle, examining gliding and host-cell invasion as distinct MyoA-dependent processes, and using biological markers to resolve defects during intracellular development in processes linked to other parasite myosins, particularly MyoF and MyoJ, including apicoplast division and inheritance, daughter-cell budding, intracellular trafficking, and late daughter-cell maturation and constriction.

PCIP is a reversible allosteric myosin inhibitor whose biochemical and structural properties were characterized using mammalian class-I myosins and several *Dictyostelium discoideum* motors. PCIP potently reduced actin-activated turnover of class-I myosins (IC₅₀ ∼5 µM), while showing substantially weaker activity against Myo2 and Myo5b and no detectable activity against human myosin-6 or myosin-7a motor-domain constructs *in vitro* (<u>Chinthalapudi et al., 2011</u>). Although a co-crystal structure of PCIP-bound *D. discoideum* Myo2 motor domain was reported (<u>Preller et al., 2011)</u>, the poor electron density for PCIP (real-space correlation coefficient, 0.39) limits confident assignment of its binding site. More recently, PCIP treatment was shown to impair *Trypanosoma cruzi* flagellar-based motility, host-cell adhesion and invasion. Pretreatment of host cells also reduced parasite invasion, probably by interfering with lysosome recruitment to the entry site, but direct engagement of a *T. cruzi* myosin was not demonstrated (<u>Spirandelli da Costa et al., 2022</u>). These observations prompted us to investigate whether PCIP could directly perturb myosin-dependent processes throughout the *T. gondii* lytic cycle.

Structural modeling with AF3 provided initial support for this possibility, with recurrent PCIP placements predicted within the motor domains of MyoA, MyoF, and MyoJ, but not MyoH. Notably, the predicted poses differed among these motors. In MyoA, PCIP consistently localized near the KNX-binding site defined by co-crystal structure, whereas recurrent placements in MyoF and MyoJ were more distant from the corresponding region. Given the limitations of AF3 ligand placement, however, these differences may still be compatible with binding to a similar site with different affinities. More generally, because the binding site was not defined *a priori*, these predictions remain exploratory and neither establish direct PCIP-myosin binding nor identify *bona fide* binding sites. Rather, they provide a structural framework for interpreting the distinct PCIP-induced phenotypes and for prioritizing candidate myosins for future biochemical and structural validation.

Guided by these structural predictions, we next used an integrated cell-biological approach to characterize PCIP-induced defects at successive stages of the tachyzoite lytic cycle, beginning with the well-established MyoA-dependent processes of gliding and invasion. Among apicomplexan myosins, MyoA remains the most extensively validated pharmacological target. The recently identified inhibitor UCB-9721 inhibits *T. gondii* motility in a 3D matrix with an IC₅₀ of 0.139 µM (<u>Snyder et al., 2026</u>), while KNX-115 inhibits *P. falciparum* sporozoite 2D gliding at approximately 0.117 µM (<u>Trivedi et al., 2026)</u>. Although differences in assay geometry, recording duration, and quantitative endpoints preclude direct comparison of potencies, PCIP impaired productive 2D gliding within similar nanomolar concentration range. These findings establish gliding motility as a particularly PCIP-sensitive process and are consistent with interference with MyoA-dependent motor activity.

Calcium-ionophore stimulation provided an additional means of challenging PCIP-induced motility arrest by strongly activating the Ca²⁺-dependent motility machinery. Productive gliding requires coordination between micronemal adhesin delivery and actomyosin-driven rearward translocation. The ability of ionophore stimulation to overcome PCIP-induced arrest may therefore reflect enhanced adhesive-cargo availability and motor activation, allowing productive movement despite partial inhibition. However, ionophores simultaneously stimulate microneme discharge, conoid protrusion, F-actin dynamics, and actomyosin activation, precluding assignment of the rescue to a single mechanism. These experiments therefore primarily demonstrate that PCIP-induced motility arrest is not irreversible and can be overcome by strong activation of the Ca²⁺-dependent motility program.

PCIP also inhibited host-cell invasion in a dose-dependent manner, although at slightly higher concentrations than those suppressing gliding, which may appear unexpected given the established contribution of the actomyosin machinery to entry (<u>Andenmatten et al., 2013</u>). However, this difference may reflect distinct thresholds of motor activity, with residual MyoA activity and/or compensatory host-cell forces supporting sub-optimal invasion when active invasion is strongly impaired (<u>Egarter et al., 2014</u>; <u>Bichet et al., 2016</u>). This interpretation is consistent with evidence that minimal gliding capacity can support invasion (<u>Bichet et al., 2014</u>), suggesting different MyoA activity requirements for these processes.

A particularly striking effect of PCIP was the concentration-dependent inhibition of intracellular growth, which became significant at 0.4–0.5 µM. By combining live-cell imaging and expansion microscopy with markers of defined events during endodyogeny, we resolved this phenotype beyond a general arrest of replication. PCIP disrupted early daughter-cell development, impairing Myo F-dependent apicoplast duplication as well as daughter IMC and cortical MT assembly.

The convergence of these cellular defects is compatible with perturbation of MyoF-dependent functions, given the established involvement of this motor early steps of endodyogeny including apicoplast inheritance, daughter-cell budding, and intracellular trafficking. Moreover, an 8-h exposure to 0.5 µM PCIP, sufficient to arrest the first round of endodyogeny, was not fully overcome following drug removal. Intracellular replication therefore represents a major and persistent point of PCIP action.

A distinct phenotype emerged at lower PCIP concentrations (0.1 µM) that still permitted daughter-cell formation. Progeny frequently lost plasma-membrane integrity before egress, releasing cytosolic GFP into the lumen of an otherwise intact PV. This phenotype was also observed independently of live imaging and increased at 0.25 µM PCIP, arguing against imaging-associated phototoxicity. Its occurrence after daughter-cell formation distinguishes it from the earlier developmental arrest observed at higher PCIP concentrations and suggests that PCIP also interferes with processes required for late daughter-cell maturation and integrity. MyoJ, MyoI, and potentially MyoE, which function in distinct compartments during late daughter-cell development and organization, represent plausible contributors to this phenotype, although their involvement remains to be established directly.

Collectively, PCIP induced a concentration- and time-dependent spectrum of defects across the *T. gondii* lytic cycle, from impaired motility and invasion to arrested intracellular development, defective daughter-cell maturation and loss of progeny integrity. The convergence of these phenotypes with structural predictions for MyoF, MyoJ and MyoA raises the possibility that PCIP interferes with multiple parasite motors, although this model remains to be demonstrated directly. Identifying PCIP targets and their binding sites will determine whether a common allosteric pocket can be exploited across functionally distinct apicomplexan myosins and which motor functions represent the most pharmacologically vulnerable points of the parasite life cycle. Extending this approach to other Apicomplexa should establish whether multi-myosin interference can complement selective MyoA inhibition as a broader antiparasitic strategy.

## Acknowledgments

### Author contributions

**Romane Agresti:** Investigation, Formal analysis, Data curation, Visualization, Methodology, writing legends, review & editing. **Faraj Gtat:** Investigation, Formal analysis, Data curation, Visualization, Methodology, writing legends, review & editing. **Guillaume Drumont:** Investigation, Formal analysis, Data curation, Visualization, Software, review and editing. **Bastien Touquet:** Investigation, Methodology, Formal analysis, Data curation, Visualization. **Christopher Swale:** Methodology, Software, Formal analysis, Visualization, Writing, review and editing. **Isabelle Tardieux:** Conceptualization, Methodology, Supervision, Project administration, Funding acquisition, Investigation, Formal analysis, Writing original draft, Writing, review and editing.

### Funding

This work was supported by the Laboratoire d’Excellence ParaFrap, France (ANR-11-LABX-0024), awarded to I.T. and used for G.D salary and required reagents and material, and by the Agence Nationale de la Recherche (ANR) grant ANR-23-CE44-0024 (MiniToxoAd; I.T., Principal Investigator), which supported R.A.’s salary.

### Disclosure AI

We used ChatGPT (OpenAI) to polish syntax and grammar throughout the text and to reorganize the initial draft. We used Claude (Anthropic) for AI-assisted coding of the macros designed for the invasion assay.

## Data availability statement

All data will be available upon request. Data relating to invasion assays analysis including Fiji macros are deposited on our Github as explained in the Methods.

## Supplemental Figure Legends

**Figure S1:**
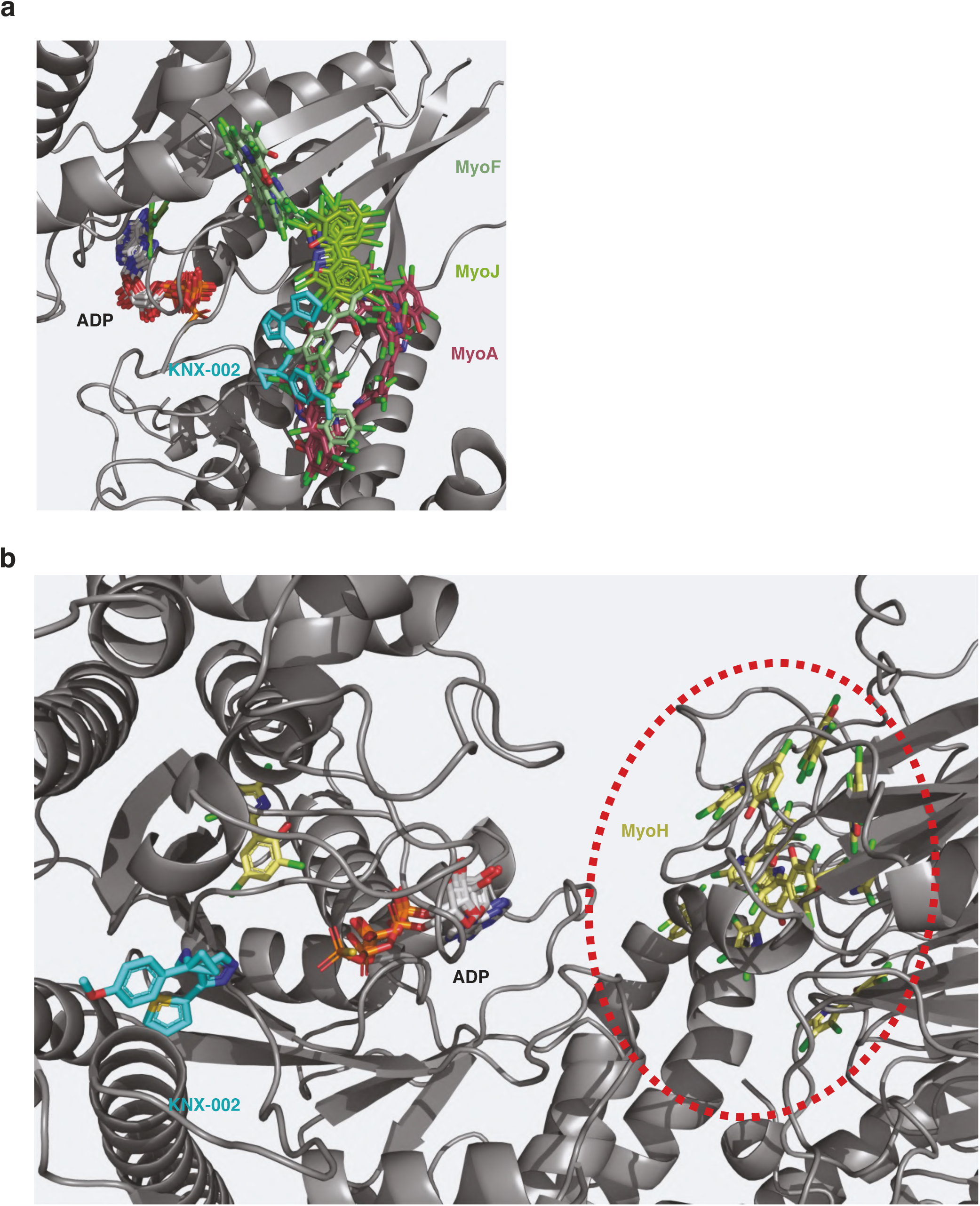
**(a)** Superposition of PCIP placements generated by AF3 co-folding as shown in Fig 1c. PCIP atoms are colored according to the CPK conventions, with carbons retaining the color associated with their respective myosin. **(b)** AF3 generated PCIP placements on the MyoH protein. The different PCIP positions (yellow) are diffused on the β-propeller of the protein, outside of the motor domain. For both figures, ADP molecules from the AF3 co-folding predictions are represented according to the CPK convention, with carbons in white. The MyoA inhibitor KNX-002 (cyan) is shown from *P. falciparum* MyoA based on structural resolution.

**Figure S3:**
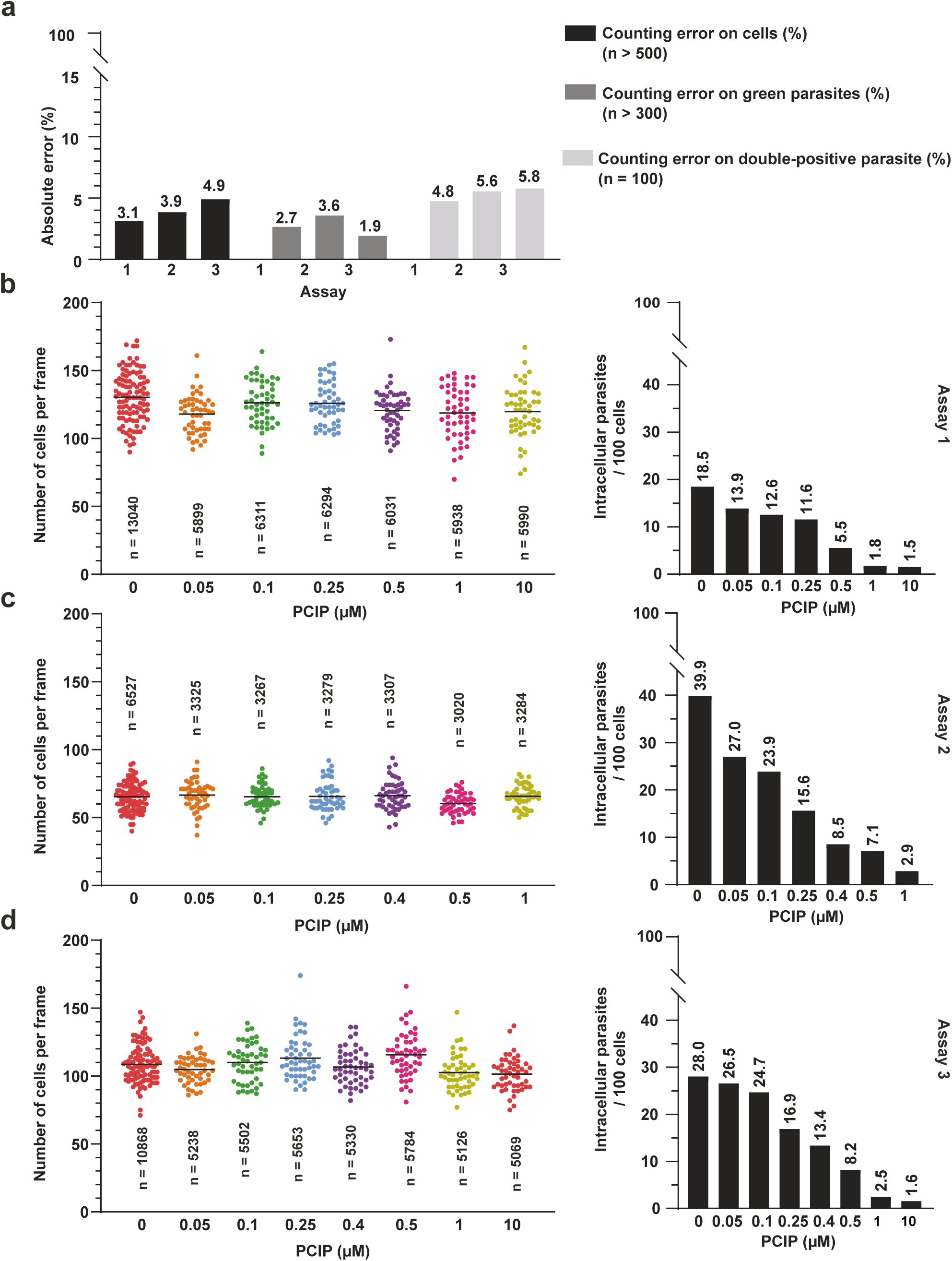
Validation and per-experiment quantification of the invasion assay. **(a)** Accuracy of automated cell and parasite counting, assessed by comparing macro-generated counts with manual counts from a randomly selected subset of images comprising approximately 500 HFF cells, 300 GFP-expressing parasites, and 100 GFP-SAG1 double-positive parasites for each of the three independent assays. Counting errors are expressed as absolute percentage errors. Using absolute values prevents overestimation and underestimation in individual images from cancelling each other out and thereby underestimating the overall error. Errors remained below 6% for all three counting categories across all assays **(b–d)** Individual data from invasion assays 1, 2, and 3, respectively. **Left:** Number of HFF cells per field across PCIP treatment conditions. Each dot represents one field. Within each experiment, 50 fields were analyzed per PCIP concentration and 100 fields for the vehicle-treated control. Horizontal lines indicate the mean, and the total number of cells analyzed for each condition is indicated. Cell density was comparable across treatment conditions within each experiment. **Right:** Invasion rates for the corresponding experiments, expressed as the number of intracellular parasites per 100 host cells. PCIP reduced parasite invasion in a dose-dependent manner in each of the three independent assays, with mixed-effects model analysis revealing a significant linear trend in all experiments (p < 0.0001). The 0.4 μM condition was not included in assay 1, whereas the 10 μM condition was not included in assay 2.

**Sup Table 1:** PCIP cytotoxicity in the immortalized human A549 epithelial cell line and in HFFs. The number of cells detected on the Hoechst positive nuclei and Sytox Green positive cells are presented for the two independent assays. n=2 for each concentration except for * where n=1 replicate.

| [PCIP]<br>(μM) | A549 cells | | | | HFF cells | | | |
| --- | --- | --- | --- | --- | --- | --- | --- | --- |
|  | Assay 1 |  | Assay 2 |  | Assay 1 |  | Assay 2 |  |
|  | DAPI +<br>(nb cells) | SYTOX +<br>(%) | DAPI +<br>(nb cells) | SYTOX +<br>(%) | DAPI +<br>(nb cells) | SYTOX +<br>(%) | DAPI +<br>(nb cells) | SYTOX +<br>(%) |
| 0 | 20782* | 0,1 | 6793* | 0,12 | 2909* | 2,6 | 5287 | 0,06 |
| 0,1 | 18880 | 0,2 | 5142 | 0,3 | 7103 | 2,0 | 5780 | 0,2 |
| 0,25 | 19865 | 0,3 | 5030* | 0,1 | 6516 | 3,0 | 5361 | 0,1 |
| 0,5 | 18236 | 0,3 | 5841 | 0,1 | 6904 | 2,0 | 4700 | 0,1 |
| 0,75 | 14795 | 0,2 | 5005* | 0,0 | 6957 | 1,8 | 4655 | 0,1 |
| 1 | 13881 | 0,1 | 4540 | 0,1 | 6512 | 2,1 | 4143 | 0,1 |
| 4 | 13133 | 0,1 | 7236* | 0,2 | 5448 | 2,0 | 3882 | 0,0 |
| 8 | 15514 | 0,1 | 2936* | 0,6 | 3971 | 2,5 | 3190 | 0,2 |
| 10 | 14882 | 0,2 | 2948 | 0,8 | 4683 | 2,2 | 3379 | 0,1 |
| 20 | 14698 | 0,2 | 4334* | 0,7 | 5179 | 2,6 | 3412 | 0,4 |
| 40 | 14229 | 0,2 | 3684* | 0,5 | 5443 | 2,3 | 3139 | 0,3 |
| 80 | 11394 | 0,1 | 3796 | 0,4 | 4023 | 3,6 | 2765 | 0,8 |
| Ctrl + | / | / | 7111* | 100,0 | 1990* | 98,6 | / | / |

## Supplementary Movie Legends

**Movie 1: Tachyzoite productive 2D motion in response to increasing PCIP concentrations.** A time-lapse DIC video showing alterations in tachyzoite gliding behavior during exposure to increasing concentrations of PCIP (from left to right): vehicle, 0.1 µM, 0.5 µM, and 1 µM. Scale bar and time are indicated in the movie. Note that typical helical and circular trajectories are observed in tachyzoites exposed to either vehicle or 0.1 µM PCIP.

**Movie 2: Recovery of tachyzoite motility following PCIP removal.** A time-lapse DIC video showing recovery of tachyzoite motility 30 min after PCIP removal. The vehicle control is shown on the left, while tachyzoites previously treated with 1 µM PCIP are shown on the right. Following PCIP removal, tachyzoites recover productive gliding motility, indicating that the inhibitory effect of PCIP on motility is reversible. Scale bar and time are indicated in the movie.

**Movie 3: Calcium ionophore A23187 restores motility in PCIP-treated tachyzoites.** A time-lapse DIC video showing tachyzoite motility following treatment with the calcium ionophore A23187. Tachyzoites treated with A23187 alone (vehicle control) are shown on the left, while tachyzoites pre-treated with 1 µM PCIP followed by A23187 treatment are shown on the right. A23187 treatment restores productive gliding motility in PCIP-treated tachyzoites. Scale bar and time are indicated in the movie.

**Movie 4: Time-lapse imaging of tachyzoite invasion.** Time-lapse DIC and fluorescence overlay imaging showing RON2-mC–labelled tachyzoites (red) invading U2OS cells expressing a GFP-GPI plasma membrane marker (green) under vehicle control (top) or PCIP treatment (bottom) conditions. Note that the main features of parasite entry, including RON2 insertion and host plasma membrane budding, are preserved under PCIP treatment, with similar characteristics and kinetics to those observed under vehicle conditions. Scale bar and time are indicated in the movie.

**Movie 5: Time-lapse imaging of vacuole growth over 28 h following parasite invasion.** Time-lapse fluorescence imaging of tachyzoites co-expressing IMC12-mC (magenta) and BCC7-Em (green) following invasion under vehicle control (top) or 0.1 µM PCIP treatment (bottom) conditions. Under both conditions, parasites undergo similar rounds of endodyogeny, characterized by maternal IMC collapse and reassembly during daughter cell budding, resulting in comparable patterns of intracellular growth over the 28-h observation period. Scale bar and time are indicated in the movie.

**Movie 6: Time-lapse imaging of vacuole growth over 21 h following parasite invasion.** Time-lapse fluorescence imaging of tachyzoites co-expressing IMC12-mC (magenta) and BCC7-Em (green) following invasion and exposure to 0.5 µM PCIP. In contrast to the vehicle condition, no detectable dynamics of IMC12 or BCC7 are observed in PCIP-treated parasites, correlating with the absence of daughter cell budding throughout the observation period. These findings suggest that PCIP interferes with an early step of endodyogeny, prior to the initiation of daughter cell budding. Scale bar and time are indicated in the movie.

**Movie 7: Time-lapse imaging of vacuole growth over 30 h following parasite invasion.** Time-lapse fluorescence imaging of tachyzoites co-expressing P12-mC (magenta) and BCC7-Em (green) following invasion under vehicle control conditions. During successive rounds of endodyogeny, the nucleus undergoes characteristic changes in shape, while BCC7 dynamically redistributes from the apical to the basal pole of maturing daughter cells. These coordinated dynamics accompany daughter cell development and maturation throughout intracellular replication. Scale bar and time are indicated in the movie.

**Movie 8: Time-lapse imaging of vacuole growth over 30 h following parasite invasion.** Time-lapse fluorescence imaging of tachyzoites co-expressing P12-mC (magenta) and BCC7-Em (green) following invasion under 0.5 µM PCIP treatment. In contrast to the vehicle control, (movie 7) no characteristic nuclear fork associated with endodyogeny is observed, while BCC7 remains localized at the basal pole of individual parasites throughout the observation period. The absence of these dynamics is consistent with a PCIP-induced block at an early step of endodyogeny. Scale bar and time are indicated in the movie.

**Movie 9: Intracellular replication, egress, and reinvasion under vehicle conditions.** Time-lapse fluorescence imaging of GFP-expressing tachyzoites under vehicle control conditions following invasion. Imaging was initiated approximately 30 h post-invasion and continued for 24 h, capturing late rounds of endodyogeny followed by parasite egress and subsequent reinvasion of neighboring host cells by the progeny. Scale bar and time are indicated in the movie.

**Movie 10: Intracellular replication and parasite lysis under 0.1 µM PCIP treatment.** Time-lapse fluorescence imaging of GFP-expressing tachyzoites continuously exposed to 0.1 µM PCIP following invasion. Imaging was initiated approximately 30 h post-invasion and continued for 24 h. In several PVs, parasites undergo lysis, as indicated by the loss of cytosolic GFP, while remaining confined within the PV, consistent with preserved PV integrity. Host cells remain morphologically intact. In the rare event of host cell rupture and progeny release, no subsequent invasion of neighboring cells is observed. Scale bar and time are indicated in the movie.

